# Seroprotection and improved polyfunctional immunity conferred by a YF17D-vectored rabies vaccine

**DOI:** 10.64898/2026.11.15.766779

**Authors:** Lorena Sanchez-Felipe, Yeranddy A. Alpizar, Sanne Terryn, Guanghui Wu, Babs E. Verstrepen, Hermann Giresse Tima, Pauline Percier-Lehebel, Gerrit Koopman, Kinga P. Böszörményi, Niraj Mishra, Sarah Debaveye, Katrien Geerts, Ji Ma, Thomas Vercruysse, Hendrik Jan Thibaut, Ashley C. Banyard, Pieter Leyssen, Corinne Vandermeulen, Willy Bogers, Olivier Denis, Marta Romano, Ernst Verschoor, Anthony R. Fooks, Steven Van Gucht, Johan Neyts, Lotte Coelmont, Kai Dallmeier

**Affiliations:** KU Leuven Department of Microbiology, Immunology and Transplantation, Rega Institute, Virology, Antiviral Drug and Vaccine Research, Laboratory of Molecular Vaccinology and Vaccine Discovery (MVVD), Leuven, Belgium; Viral Diseases-Infectious Diseases in Humans, Sciensano, Brussels, Belgium; Animal and Plant Health Agency (APHA), Department of Virology, Weybridge, New Haw, Surrey, KT15 3NB, UK; Department of Virology, Biomedical Primate Research Centre (BPRC), Rijswijk, The Netherlands; Immune Response-Infectious Diseases in Humans, Sciensano, Brussels, Belgium; KU Leuven Department of Microbiology, Immunology & Transplantation, Rega Institute, Virology, Antiviral Drug & Vaccine Research, Laboratory of Virology & Antiviral Research, Leuven, Belgium; KU Leuven Department of Microbiology, Immunology & Transplantation, Rega Institute, Translational Platform Virus, Vaccine and Cancer Research (TPVC), Leuven, Belgium; KU Leuven Department of Public Health and Primary Care, Leuven University Vaccinology Center, Leuven, Belgium; Centre of Expertise Perspective in Health, Avans Hogeschool, Breda, Netherlands; GSK, Wavre, Belgium; InhaTarget Therapeutics, Gosselies, Belgium; Aurigene Oncology, Hyderabad, India; Army Medical University, ChongQing, China; Astrivax Therapeutics NV, Heverlee, Belgium

## Abstract

Rabies remains a major public-health challenge, with pre-exposure vaccination strategies constrained by the need for multiple doses and repeated healthcare visits. Here we describe YF-RabG, a live-attenuated rabies vaccine candidate based on the yellow fever 17D (YF17D) vaccine backbone that stably expresses the rabies virus glycoprotein (RabG). YF-RabG exhibited an attenuated phenotype and favourable safety profile compared with parental YF17D across several preclinical models, including neonatal and immunocompromised mice. A single immunization induced robust humoral and cell-mediated immune responses in non-human primates. Rabies virus-neutralizing antibody titres readily exceeded the widely accepted correlate of protection in mice, hamsters and non-human primates by several orders. Accordingly, vaccination conferred protection against lethal rabies challenge in a range of stringent mouse models, maintained seroprotective antibody levels for at least one year, and increased following a boost using a licensed rabies vaccine. Furthermore, the YF17D vector was readily adapted to accommodate glycoproteins from distant lyssaviruses, generating protective vaccine candidates against Mokola, Lagos bat and Lleida bat lyssaviruses. Collectively, these findings identify YF-RabG as a promising single-dose rabies vaccine candidate and highlight the versatility of the YF17D platform for lyssavirus vaccine development.

## INTRODUCTION

Rabies is a severe and almost uniformly fatal, yet widely neglected, zoonotic disease caused by viruses belonging to the *Lyssavirus* genus^1–3^. Most cases affecting wildlife, domestic animals, and humans, are attributed to the prototypic rabies virus (RABV). Human cases are predominantly due to dog-bites transmitted RABV, accounting for over 99% of reported incidents. Despite efforts to control the disease at its source through culling or immunizing stray dogs, at least 59,000 human deaths occur annually, with 40% of these victims being children under 15 years of age^1,4^. Many cases are believed to go unreported, particularly in regions where the disease is prevalent, namely Africa and Asia^5,6^.

Human rabies is entirely preventable through vaccination. Current vaccines consist of whole inactivated virus, typically cultured on chicken embryo cells (e.g. Rabipur^®^ by GSK) or mammalian cell lines of monkey origin^7^. They can be used for both pre-exposure prophylaxis (PrEP) for individuals at risk of exposure and post-exposure prophylaxis (PEP) for those who may have been exposed to lyssaviruses. In unvaccinated individuals, PEP may include rabies-specific immunoglobulins (RIG) in addition to vaccination, depending on the nature of the exposure and applicable guidelines^8,9^. However, PEP must be administered promptly and completed according to the appropriate vaccination schedule, which in previously unvaccinated individuals commonly consists of a 4-dose regimen, to effectively interrupt the otherwise almost invariably fatal progression to rabies encephalitis. Unfortunately, the complexity of the multi-dose vaccine regimen and shortage of vaccines leave many in urgent need unprotected and untreated. Likewise, while the use of current multi-dose vaccines for PrEP in children at risk living in highly endemic areas has proven effective in reducing the disease burden^10–13^, population-wide roll-out is hindered by prohibitively high costs^2,14^. Hence, there is a need for next-generation rabies vaccines with improved immunogenicity, sufficient to confer long-term protection following a single-dose administration that can be included in routine childhood immunization programs to broaden global PrEP coverage^15,16^.

The yellow fever 17D (YF17D) vaccine, developed in the 1930s, is a live-attenuated vaccine that has proven to be highly effective, providing long-lasting, possibly lifelong, immunity from a single dose^17^. Due to its strong immunogenicity and safety profile, the YF17D vaccine has been explored as a vector for developing new vaccines against other diseases^18–23^. We and others have successfully inserted genes from various pathogens into the YF17D genome creating recombinant vaccines that can induce protective immune responses against multiple diseases at once. Use of YF17D as a vaccine vector offers a cost-effective and efficient platform for developing new vaccines, especially for use in underdeveloped regions where access to multiple vaccinations remains challenging^24^.

Here, we developed YF-RabG, a YF17D-vectored rabies vaccine candidate expressing the rabies virus glycoprotein (RabG). We investigated its safety, immunogenicity and protective efficacy in complementary animal models and assessed the broader potential of the YF17D platform for vaccination against antigenically diverse lyssaviruses.

## RESULTS

### Generation and characterization of a YF-vectored rabies vaccine candidate

A live-attenuated YF-RabG vaccine candidate was generated by inserting the coding sequence of the rabies virus glycoprotein G from the Pitman-Moore strain (RabG) as an in-frame translational fusion into the E/NS1 intergenic region of an infectious cDNA clone of YF17D^24^ (**Extended Data Fig. 1a**). Transfection of the construct into BHK-21 cells yielded infectious viral progeny. Compared with parental YF17D, YF-RabG displayed a smaller plaque phenotype and slightly reduced replication kinetics, indicative of additional replicative attenuation conferred by insertion of the heterologous antigen (**Extended Data Fig. 1b**).

As insertion and expression of foreign antigens can compromise the genetic stability of live viral vectors, we next assessed maintenance of the RabG transgene during serial passaging. RT-PCR fingerprinting confirmed preservation of the inserted RabG sequence over at least 11 consecutive *in vitro* passages, demonstrating genetic stability of the recombinant virus during prolonged replication (**Extended Data Fig. 1c**).

Expression of both vector- and transgene-derived antigens was confirmed by immunofluorescence analysis of YF-RabG-infected BHK-21 cells. Following permeabilization, YF17D and RabG antigens were intracellularly detected using polyclonal anti-YF17D serum and a RabG-specific monoclonal antibody, respectively (**Extended Data Fig. 1d**). Surface expression of RabG was further demonstrated by immunostaining of non-permeabilized cells, confirming proper trafficking of the glycoprotein to the plasma membrane of YF-RabG transduced cells (**Extended Data Fig. 1d**).

To verify that expression of RabG did not alter the structural composition of the viral particle and to exclude any risk of a change in tropism by pseudotyping with a neurotropic envelope, YF-RabG virus particles were subjected to plaque reduction neutralization assays (PRNTs) using high-titred anti-YFV and anti-RABV immune sera. YF-RabG was readily neutralized by sera from Stamaril^®^-vaccinated animals, whereas sera from Rabipur^®^-vaccinated animals failed to reduce plaque formation (**Extended Data Fig. 1e,f**). Neutralization by anti-YFV sera remained evident across serial dilutions and was still detectable at a dilution of 1:6,600. These data confirm that YF-RabG particles retain the envelope architecture and antigenic identity of genuine YF17D and do not acquire a rabies glycoprotein-coated chimeric surface phenotype.

To further assess the impact of RabG insertion on vaccine safety, we evaluated neurovirulence in neonatal BALB/c mice. Intracranial inoculation of YF-RabG resulted in significantly reduced neurovirulence compared with parental YF17D. Survival of three-to five-day-old pups inoculated with 1,000 PFU of YF-RabG (n = 11; median time to humane endpoint, 8 days; IQR, 7–8 days) was prolonged relative to animals receiving 10 PFU of YF17D (n = 11; median time to humane endpoint, 6 days; IQR, 6–6 days; P < 0.0001, log-rank test) (**Extended Data Fig. 1g,h**). Importantly, residual neurovirulence remained detectable following intracranial inoculation, consistent with retained replication competence *in vivo*, a defining feature of live-attenuated vaccines. This observation is in line with the historical use of neonatal mouse neurovirulence assays for potency testing of YF17D-derived vaccines^25^.

### Immunogenicity of YF-RabG in non-human primates

To evaluate the immunogenicity of YF-RabG in a translationally relevant step-up model, cynomolgus macaques were vaccinated with YF-RabG and compared directly with animals receiving the licensed rabies and yellow fever vaccines Rabipur^®^ and Stamaril^®^, respectively (**Fig. 1a**). Six macaques received YF-RabG (10^5^ PFU; s.c.)^20^, whereas a control group (n = 6) received Rabipur^®^ (one human dose at days −7 and 0; i.m.) together with Stamaril^®^ (one human dose at day 0; s.c.). All animals received a homologous booster immunization at day 28. The vaccines were well tolerated, with no treatment-related changes in clinical observations, body weight, body temperature, injection site reactions, clinical chemistry or hematological parameters (**Extended Data Figs. 2 and 3**).

**Fig. 1:**
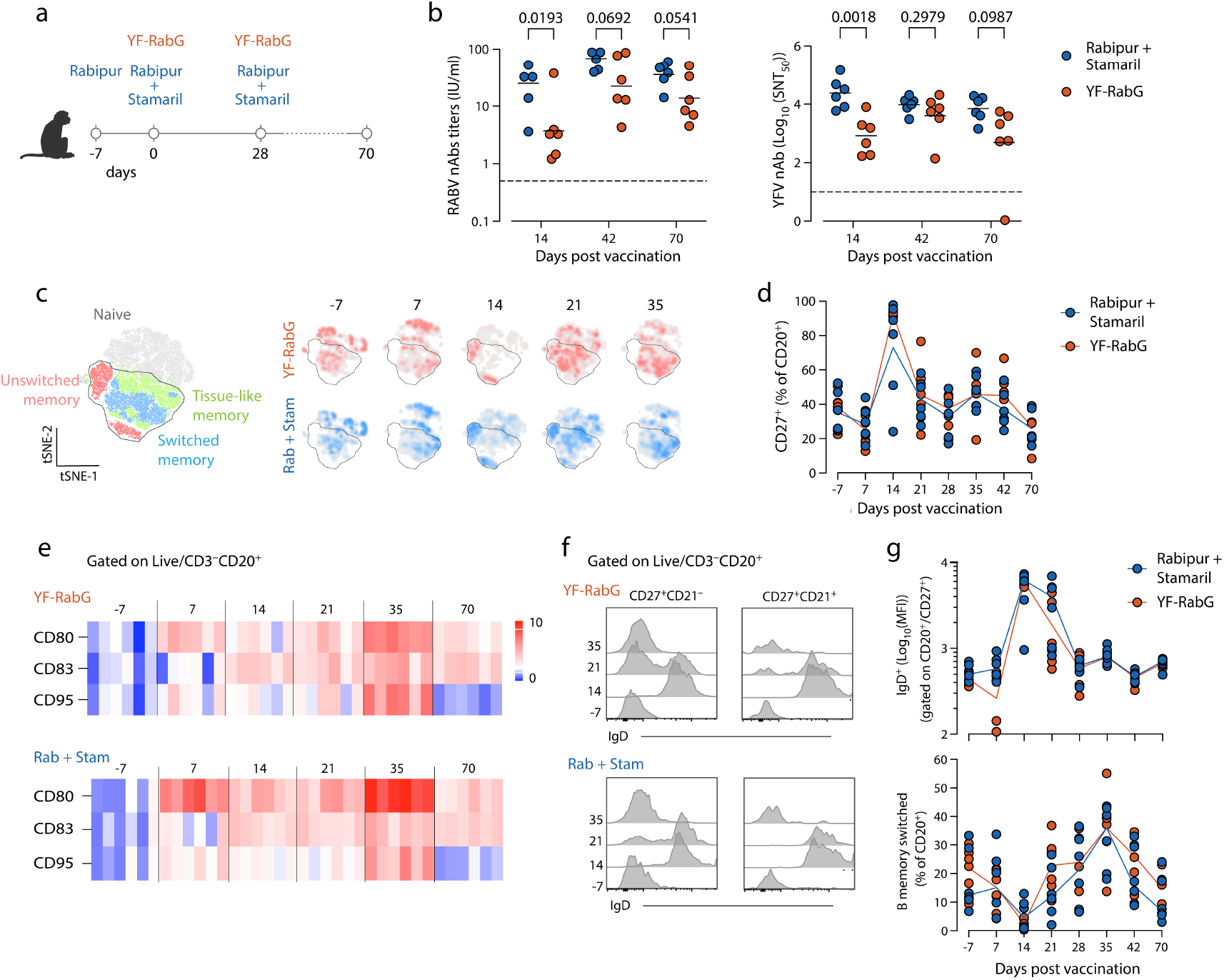
Immunogenicity of YF-RabG vaccine candidate in NHP. a, Vaccination schedule in Cynomolgus macaques (n = 6 per group). Animals received either YF-RabG (10^5^ PFU at days 0 and 28) or a prime dose with Rabipur^®^ (day -7), followed by simultaneous vaccinations with Rabipur^®^ and Stamaril^®^ (days 0 and 28). **b,** RABV- and YFV-neutralizing antibody titers at days 14, 42 and 70. Data points represent individual animals; bars indicate geometric (left) and arithmetic (right) mean values. Statistical significance was assessed using mixed-effects model fitted by restricted maximum likelihood (REML), with Geisser–Greenhouse correction, followed by two-sided Šídák’s multiple-comparisons test. The dotted line indicates the correlate of protection threshold (RABV nAbs, 0.5 IU/mL; YFV nAbs, 1 Log_10_(SNT_50_)). **c,** Left, t-distributed stochastic neighbour embedding (t-SNE) map of CD20^+^ cells showing phenotypically distinct naïve and memory B-cell clusters. The outlined region encompasses the entire memory B-cell population. Right, density distributions across the t-SNE map at different time points for the YF-RabG (red) and Rabipur^®^+Stamaril^®^ (Rab+Stam, blue) groups. **d**, Frequency of circulating memory B cells (CD27^+^ cells gated within the CD20^+^ population) at different time points after vaccination. Lines indicate group means; dots represent individual animals. **e**, Heatmap showing the expression of the activation markers CD80, CD83 and CD95 at different time points after vaccination. Values are expressed as z scores relative to the baseline measurement on day −7. **f**, IgD expression in circulating resting (CD27^+^CD21^+^) and activated (CD27^+^CD21^−^) memory B-cell populations on days −7, 14, 21 and 35. **g**, Expression levels of IgD in circulating memory B cells (top) and frequency of class-switched (IgD^−^) memory B cells (bottom), expressed as percentages of the total CD20^+^ B-cell population. Lines indicate group means; dots represent individual animals.

A single immunization with YF-RabG elicited robust RABV-neutralizing antibody (nAbs) responses in all animals. By day 14, all macaques had seroconverted and reached antibody levels above the accepted CoP (0.5 IU/ml), despite receiving only a single dose of YF-RabG compared with two administrations of Rabipur^®^ in the control group (**Fig. 1b**). RABV-nAb titres were lower in YF-RabG-immunized animals than in the twice Rabipur^®^/Stamaril^®^ group at day 14 (GMT [95% CI]: 3.7 [1.0-15.4] *vs* 25.7 [7.5-88.0], respectively; *P =* 0.0193). Following booster immunization, RABV-neutralizing antibody titres increased further in the YF-RabG group, reaching high GMTs of 22.9 IU/ml (95% CI, 6.8-76.9) at day 42, indicating efficient recall of vaccine-induced immunity (**Fig. 1b; Extended Data Fig. 4a**).

As expected, YF-RabG also induced potent immunity against YFV. All animals developed YFV-neutralizing antibody titres exceeding the FDA-defined surrogate threshold for protection (Log_10_ neutralization index, LNI > 0.7) after a single immunization^26^.

Although, at day 14, YFV-neutralizing antibody titres were initially lower in YF-RabG-vaccinated macaques than in animals receiving Stamaril^®^ (mean Log10 nAb titre ± SD: 3.0 ± 0.7 *vs* 4.4 ± 0.5; *P =* 0.0018), titres increased following booster immunization and became comparable to those induced by two doses of Stamaril^®^ at later time points (**Fig. 1b; Extended Data Fig. 4b**). Together, these findings demonstrate that YF-RabG induces simultaneous humoral immunity against both rabies virus and yellow fever virus in non-human primates.

Circulating immune cell dynamics were characterized by high-dimensional spectral flow cytometry. T cells (CD3, CD4 and CD8), B cells (CD20 and HLA-DR), monocytes (CD14 and CD16) and dendritic cells (CD1c and CD11c) were identified using established lineage-specific surface markers and visualized by t-distributed stochastic neighbor embedding (t-SNE) and FlowSOM clustering (**Extended Data Fig. 5**). The overall frequencies of circulating CD20^+^ B cells, CD4^+^ T cells and CD8^+^ T cells remained largely unchanged following vaccination with either YF-RabG or the Rabipur^®^/Stamaril^®^ combination (**Extended Data Fig. 5e**).

To further characterize vaccine-induced B-cell responses, CD20^+^ B cells were classified as naïve (CD27^−^CD21^+^), switched memory (CD27^+^IgD^−^), unswitched memory (CD27^+^IgD^+^) and tissue-like memory (CD27^−^CD21^−^) subsets^27^. A single dose of YF-RabG induced a rapid increase in the frequency of circulating CD27^+^ memory B cells that peaked already 14 days after vaccination (**Fig. 1d,f**; *P* < 0.0001). This early response was predominantly driven by unswitched memory B cells (CD27^+^IgD^+^), which declined by day 28 and were not further expanded following booster immunization (**Fig. 1f,g**). In contrast, administration of the second YF-RabG dose induced a marked increase in switched memory B cells (CD27^+^IgD^−^), consistent with maturation of the B-cell response (**Fig. 1f,g**). Similar dynamics were observed in animals receiving the Rabipur^®^/Stamaril^®^ combination. In both groups, expression of co-stimulatory molecules increased rapidly following vaccination and remained elevated throughout the study period (**Fig. 1e**).

We subsequently assessed memory T-cell responses. YF-RabG vaccination induced an early increase in the frequency of circulating central memory CD4^+^ T cells (T_CM_), detectable by day 7 and accompanied by increased expression of activation markers (**Extended Data Fig. 6a,b,e**). Although the frequency of effector memory CD4^+^ T cells (T_EM_) remained relatively stable (**Extended Data Fig. 6b**), this population also displayed enhanced activation following primary vaccination (**Extended Data Fig. 6e**). Likewise, both central memory and effector memory CD8^+^ T-cell populations exhibited increased activation and expansion after vaccination (**Extended Data Fig. 6c-f**).

Booster immunization at day 28 further increased the frequency of circulating CD4^+^ T_CM_ cells and was associated with prominent upregulation of the transferrin receptor CD71, a marker of T-cell activation and proliferation^28^ (**Extended Data Fig. 6b,e**). Similarly, the frequencies of both CD8^+^ T_CM_ and T_EM_ cells increased following booster vaccination (**Extended Data Fig. 6c-f**). Animals receiving Rabipur^®^/Stamaril^®^ displayed broadly comparable activation profiles, although changes in circulating CD4^+^ and CD8^+^ T_CM_ frequencies following primary vaccination were less pronounced than those observed after YF-RabG vaccination (**Extended Data Fig. 6b,d**). Nevertheless, booster immunization induced robust activation and expansion of memory T-cell populations in both groups. Together, these findings demonstrate that YF-RabG elicits coordinated B-cell and T-cell responses that accompany the development of humoral immunity in non-human primates. Rabipur^®^ (in total 3 doses) was always coadministered with Stamaril^®^ (2 doses). Therefore, the specific influence of either modality (inactivated Rabipur^®^ *vs* live Stamaril^®^) on circulating immune cell dynamics could not be differentiated.

### RABV-specific immunogenicity of YF-RabG and protection against lethal RABV challenge in mice

To evaluate whether the humoral and cellular immune responses observed in non-human primates translated into protective efficacy, YF-RabG was assessed in three murine challenge models representing increasingly restrictive conditions for YF17D replication (**Fig. 2**).

**Fig. 2:**
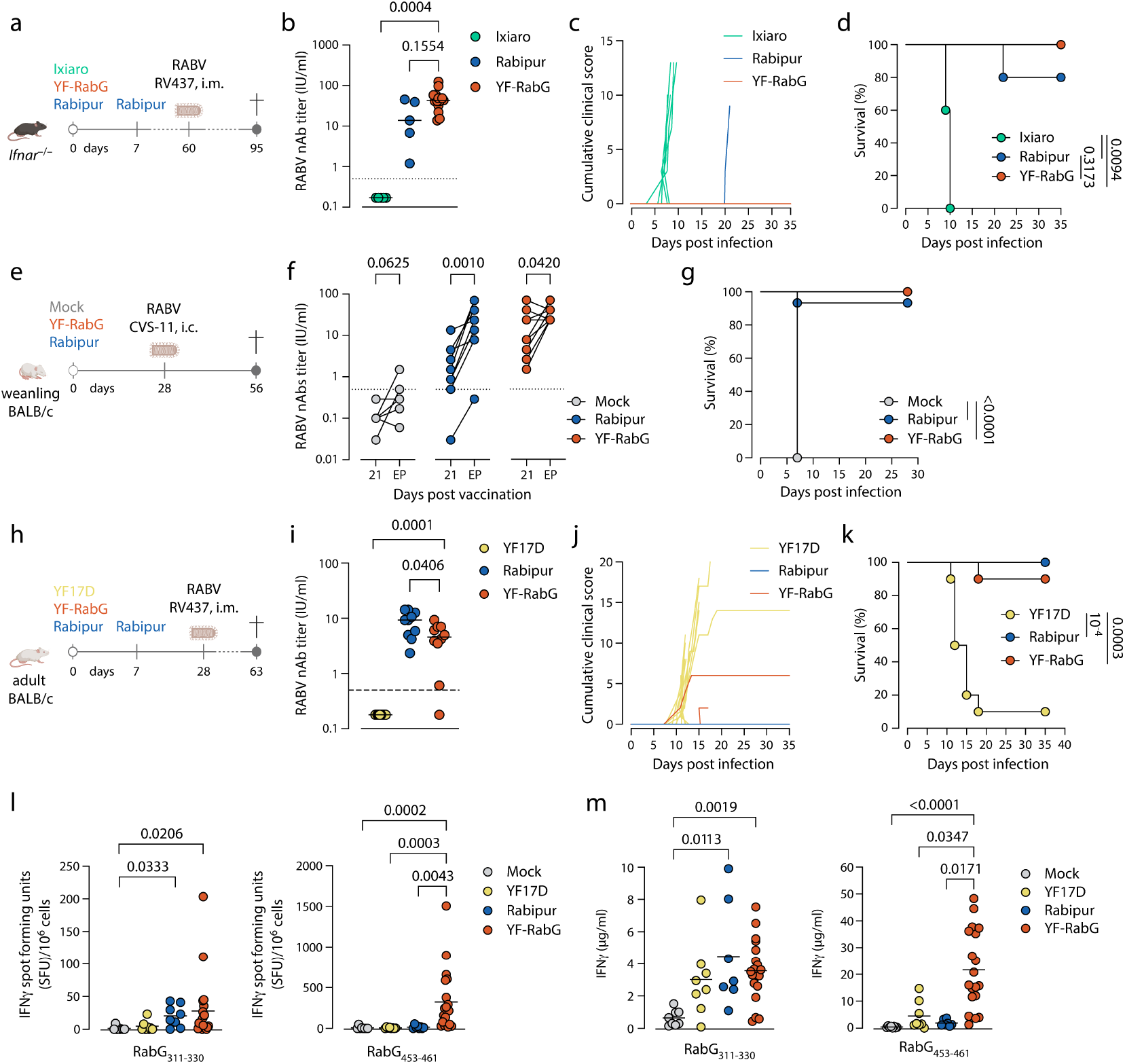
Protective efficacy of YF-RabG against RABV challenge in mice. a, Experimental workflow in *Ifnar*^−/−^ mice (6-8 weeks old). Animals received a single dose of YF-RabG (10^3^ PFU, i.p.; n = 15). Control animals received Rabipur^®^ (1/40 human dose on days 0 and 7; n = 5) or Ixiaro^®^ as an irrelevant inactivated vaccine control (day 0; n = 5). Mice were challenged intramuscularly on day 60 with RABV isolate RV437 (720 TCID_50_). **b,** RABV-neutralizing antibody titres on day 28 after vaccination. Dots represent individual animals; bars indicate means. The dotted line indicates the correlate of protection threshold (0.5 IU/ml). Two-sided Dunn’s multiple-comparisons test. **c,d,** Mean cumulative clinical score (c) and survival (d) after RABV challenge. Log-rank (Mantel–Cox) test. **e**, Experimental workflow in weanling BALB/c mice (3-4 weeks old). Animals received a single dose of YF-RabG (10^4^ PFU; n = 16), Rabipur^®^ (1/40 human dose; n = 16) or mock vaccine (2% FBS in MEM; n = 8). Mice receiving YF-RabG were administered an IFNAR1-blocking antibody (MAR1-5A3, 2 mg, i.p.) 4 h before vaccination. On day 28, mice were challenged intracranially with RABV (strain CVS-11, 50 FFU). **f**, RABV-neutralizing antibody titres measured by mFAVN on day 21 and at the humane or experimental endpoint (EP) after challenge. Dots represent individual animals; bars indicate means. The dotted line indicates the correlate of protection threshold (0.5 IU/ml). Wilcoxon matched-pairs signed-rank test. **g**, Survival after RABV challenge. Log-rank (Mantel–Cox) test. **h,** Experimental workflow in adult wild-type BALB/c mice (6–8 weeks old). Animals received a single dose of YF-RabG (2x10^3^ PFU; n = 10) or YF17D (2x10^3^ PFU; n = 10) on day 0, or Rabipur^®^ (1/40 human dose; n = 10) on days 0 and 7. Mice receiving YF-RabG or YF17D were administered an IFNAR1-blocking antibody (MAR1-5A3, 2 mg, i.p.) 4 h before vaccination. Mice were challenged intramuscularly with RABV isolate RV437 (720 TCID_50_). **i,** RABV-neutralizing antibody titres measured by RFFIT on day 28. Dots represent individual animals; bars indicate means. The dotted line indicates the correlate of protection threshold (0.5 IU/ml). Two-sided Dunn’s multiple-comparisons test. **j,k,** Mean cumulative clinical score (j) and survival (k) after RABV challenge. Log-rank (Mantel–Cox) test. **l, m,** Number of IFN-γ-secreting cells (l) and concentration of IFN-γ in culture supernatants (m) following *ex vivo* stimulation of splenocytes with the RabG-derived MHC class I and II-restricted RabG peptides RabG_453-461_ or RabG_311-330_, respectively. Splenocytes were isolated from BALB/c mice vaccinated as described in h. YF-RabG (n = 20 in l; n = 19 in m); YF17D (n = 8); Rabipur^®^ (n = 8 in l; n = 7 in m); mock-vaccinated mice (medium containing 2% FBS; n = 8). Dots represent individual animals; bars indicate means. Statistical significance was assessed using Dunn’s multiple-comparisons test; only *P* < 0.05 is shown

We first evaluated vaccine efficacy in *Ifnar1*^−/−^ mice, a fully permissive model supporting replication and hence immunogenicity of YF17D and YF17D-based vaccines^29^. Animals received a single dose of YF-RabG (10^3^ PFU), whereas control groups received either two doses of Rabipur^®^ or the unrelated inactivated Japanese encephalitis virus vaccine Ixiaro^®^ (**Fig. 2a**). By day 28, all YF-RabG-vaccinated animals had seroconverted, generating robust RABV-neutralizing antibody titres that exceeded those induced by Rabipur^®^ (GMT [95% CI]: 42.0 IU/ml [30.2-58.3] *vs* 11.54 IU/ml [1.8-73.1]; **Fig. 2b**). Following intramuscular challenge with RABV street virus isolate RV437 at day 60, all YF-RabG-vaccinated mice survived without signs of disease (15/15; 100% survival), whereas all Ixiaro^®^-vaccinated animals succumbed to infection, and one Rabipur^®^-vaccinated animal reached humane endpoints (**Fig. 2c,d**). These findings demonstrate that a single low dose of YF-RabG can be sufficient to induce complete protection from rabies in a fully permissive model. Notably, while deficiency of antiviral and immunomodulatory interferon signalling in *Ifnar1*^−/−^ mice rendered the animal highly susceptible to neurotropic RABV infection, it did not compromise live YF-RabG vaccine efficacy even from a single low dose (equivalent of ∼1/100 human dose of current YF17D vaccines).

We next assessed vaccine efficacy in immunocompetent weanling BALB/c mice, which are expected to be less restrictive to YF17D replication than adult animals^30^. To further facilitate replication of the YF17D-based vaccine vector, animals receiving YF-RabG were treated with the IFNAR1-blocking antibody MAR1-5A3 prior to vaccination^29^. Mice were vaccinated with a single dose of YF-RabG (10^4^ PFU), Rabipur^®^ or mock vaccine (2% FBS in MEM) and were challenged 28 days later with RABV strain CVS-11 via the stringent intracranial route (**Fig. 2e**). Prior to challenge, all YF-RabG-vaccinated animals seroconverted and developed RABV-neutralizing antibody titres above the CoP (GMT [95% CI]: 13.0 IU/ml [7.1-23.8]; **Fig. 2f**). Following challenge, complete protection was observed in the YF-RabG group (16/16; 100% survival), whereas 15 of 16 Rabipur^®^-vaccinated animals survived (94% survival). In contrast, all mock-immunized animals rapidly developed severe disease and reached humane endpoints (0/8 survival) (**Fig. 2g**). Notably, YF-RabG-vaccinated animals exhibited a significant increase in RABV-neutralizing antibody titres following challenge (GMT [95% CI]: 13.0 IU/ml [7.1-23.8] pre-challenge *vs* 39.0 IU/ml [27.0-56.6] at the experimental endpoint; *P =* 0.042), although several animals showed little or no increase in titre despite surviving challenge (**Fig. 2f**). A similar post-challenge boosting effect was observed in Rabipur^®^-vaccinated mice (GMT [95% CI]: 1.3 IU/ml [0.5-3.4] *vs* 23.4 IU/ml [9.5-57.8]; *P =* 0.001), consistent with an anamnestic antibody response following exposure to challenge virus. In line with this, the limited or absent boosting observed in a subset of YF-RabG-vaccinated animals, which generally displayed the highest pre-challenge RABV-neutralizing antibody titres, suggests a more effective control of challenge virus replication.

To determine whether protection could also be achieved under conditions that are less favorable for YF17D replication, we evaluated YF-RabG in adult immunocompetent BALB/c mice. Compared with the weanling BALB/c model (**Fig. 2e**), animals received a five-fold lower vaccine dose (2x10^3^ PFU) which, despite transient IFNAR blockade during vaccination, represent a substantially less permissive host environment for replication of YF17D-based vaccines^29^ (**Fig. 2h**).

Despite these restrictive conditions, a single dose of YF-RabG induced seroconversion in 90% of animals and generated RABV-neutralizing antibody titres well above the accepted CoP (GMT [95% CI]: 3.1 IU/ml [1.3-7.7]; **Fig. 2i**). Following intramuscular challenge with RABV isolate RV437, all seropositive YF-RabG-vaccinated mice survived without signs of disease, whereas nearly all YF17D-vaccinated control animals developed progressive disease and reached humane endpoints (**Fig. 2j,k**). Importantly, animals that failed to survive challenge lacked detectable RABV-neutralizing antibodies before challenge, further supporting the central role of vaccine-induced humoral immunity for protection.

To further characterize vaccine-induced cellular immunity in this stringent model, splenocytes from vaccinated mice were stimulated *ex vivo* with RabG-derived MHC class I (RabG_453–461_) and MHC class II (RabG_311–330)_ peptides. YF-RabG vaccination induced significantly higher numbers of IFN-γ-secreting cells and increased IFN-γ production compared with YF17D and mock-vaccinated controls (**Fig. 2l,m**), demonstrating the induction of RabG-specific cellular immune responses. MHC class I restricted CD8^+^ responses induced by YF-RabG appeared to exceed those for Rabipur^®^, which showed very low, if any, induction of RabG-specific CD8^+^ T cells (*P* = 0.0043, **Fig. 2l**).

Together, these results demonstrate that a single immunization with YF-RabG elicits protective immunity against rabies across murine models spanning a gradient of increasingly restrictive conditions for YF17D replication. Even under suboptimal conditions for vaccine replication, YF-RabG induced RABV-neutralizing antibody titres above the CoP and conferred robust protection against vigorous RABV challenge.

### Longevity and boostability of seroprotective RABV nAb responses

RABV-neutralizing antibody titres remained above the CoP (0.5 IU/ml) at 90 days after vaccination in all those BALB/c mice that had seroconverted following a single dose of YF-RabG (GMT [95% CI]: 3.7 [1.1-12.4] at day 90 *vs* 2.5 [0.8-7.4] at day 28; **Extended Data Fig. 7a**). These titres were comparable to those observed in animals vaccinated with 2 doses of Rabipur^®^ (*P =* 0.415 at day 28; *P =* 0.332 at day 90). Upon intramuscular challenge with RABV isolate RV437 at day 90, all animals with pre-challenge antibody titres above 0.5 IU/ml were protected (**Extended Data Fig. 7b,c**). The YF-RabG-vaccinated animals (2/10) that succumbed to challenge failed to seroconvert and lacked detectable seroprotective RABV-neutralizing antibodies throughout the study. Together, these findings demonstrate that a single dose of YF-RabG can confer durable protection against rabies for at least three months after vaccination.

To evaluate long-term immune durability in a fully immunocompetent and naturally permissive host, we next studied Syrian golden hamsters. Hamsters received a single dose of YF-RabG (either 10^3^, 5x10^3^ or 25x10^3^ PFU) or YF17D (5x10^3^ PFU) and were followed for one year (**Fig. 3a**). Three weeks after vaccination, all YF-RabG-vaccinated animals had developed RABV-neutralizing antibody titres above the accepted CoP, irrespective of the vaccine dose administered (**Fig. 3b**). Antibody titres were dose-dependent at this early time point, with the highest dose (25x10^3^ PFU) inducing significantly higher titres than the lowest dose (10^3^ PFU) (GMT [95% CI]: 18.46 [12.3-27.7] *vs* 7.0 [4.3-11.4]; *P =* 0.0282). Antibody titres gradually declined over time but remained above the CoP throughout the 52-week follow-up period. By 6 and 12 months after vaccination, no significant differences in RABV-neutralizing antibody titres were observed between dose groups, indicating long-term maintenance of seroprotective immunity irrespective of the initial vaccine dose (**Fig. 3b**).

**Fig. 3:**
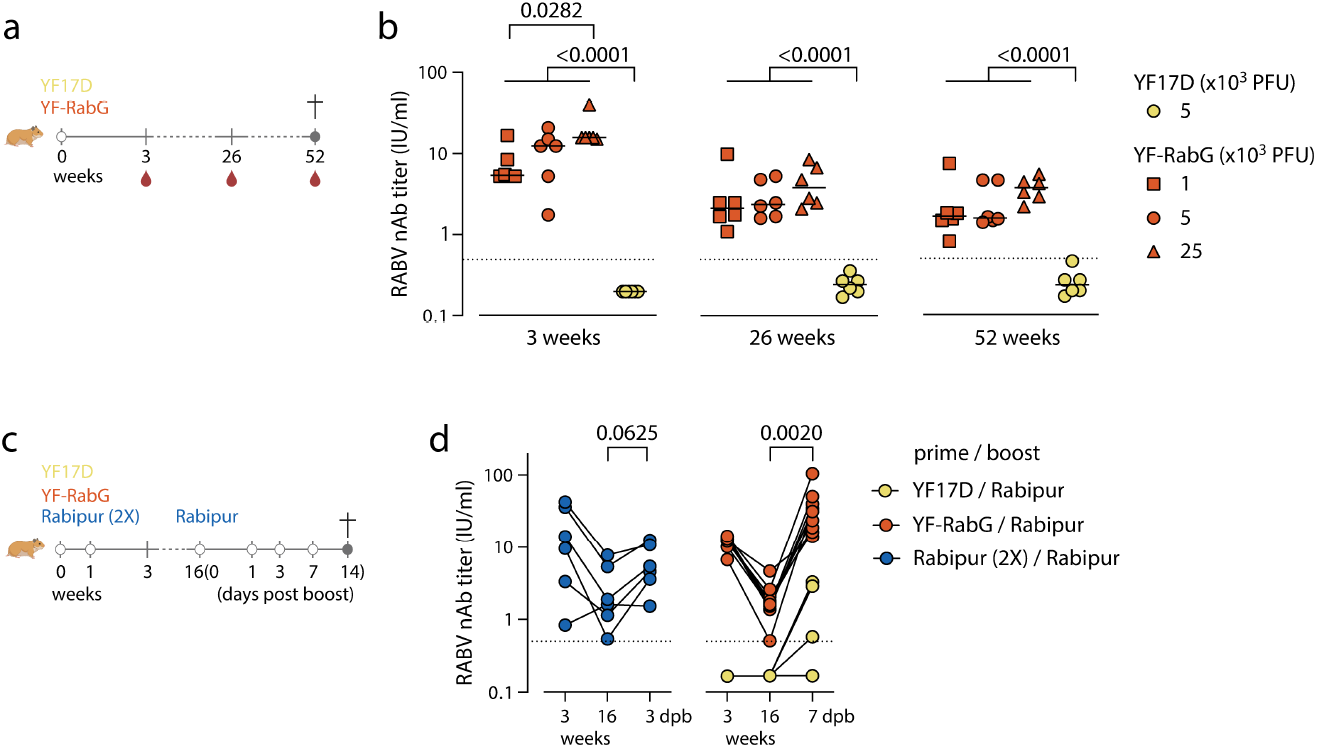
Longevity of the humoral immune responses and boostability after vaccination with YF-RabG. **a**, Experimental workflow for assessing response longevity in wild-type Syrian golden hamsters. Animals received a single dose of YF-RabG (10^3^, 5x10^3^ or 25x10^3^ PFU; n = 6 per group) or YF17D (5x10^3^ PFU, n = 6). Blood samples were collected at 3, 26 and 52 weeks after vaccination. **b**, RABV-neutralizing antibody titres at 3, 26 and 52 weeks after vaccination. Dots represent individual animals; bars indicate means. The dotted line indicates the correlate of protection threshold (0.5 IU/ml). One-way ANOVA with Tukey’s multiple-comparisons test; only *P* < 0.05 is shown. **c**, Experimental workflow for assessing response boostability in wild-type Syrian golden hamsters. Animals received YF-RabG (5x10^3^ PFU; n = 10) or YF17D (5x10^3^ PFU; n = 6) on day 0, or Rabipur^®^ (1/40 human dose; n = 6) on days 0 and 7. At week 16, all animals received a Rabipur^®^ booster (1/40 human dose). Blood samples were collected on days 1, 3, 7 and 14 after boosting. **d**, RABV-neutralizing antibody titres at 3 and 16 weeks after prime vaccination and at the peak recall response after boosting (Rabipur^®^, 3 days post boost, dpb; YF-RabG, 7 dpb). Dots represent individual animals. The dotted line indicates the correlate of protection threshold (0.5 IU/ml). One-way ANOVA with Tukey’s multiple-comparisons test; only *P* < 0.05 is shown.

For rabies prophylaxis, long-term protection relies on the ability to rapidly recall nAb responses following booster vaccination or virus exposure. We therefore assessed the boostability of YF-RabG-induced immunity in hamsters. Animals received either a single dose of YF-RabG (5x10^3^ PFU), two doses of Rabipur^®^ or YF17D (5x10^3^ PFU) as vector control (**Fig. 3c**). Consistent with the previous longevity study, YF-RabG induced RABV-nAb titres comparable to those elicited by two doses of Rabipur^®^ three weeks after vaccination (**Fig. 3d**). By week 16, antibody titres had declined from their peak values but remained above the CoP in all animals. Administration of a Rabipur^®^ booster at this time elicited a rapid and robust recall response in YF-RabG-primed animals, resulting in a rapid and marked increase in RABV-nAb titres that steadily increased until a peak at 7 days after boosting (GMT [95% CI]: 1.7 [1.1-2.5] at week 16 *vs* 30.2 [19.8-46.0] at day 7 post-boost; **Fig. 3d; Extended Data Fig. 7d-f**). A recall response was likewise observed in Rabipur^®^-vaccinated animals, although peak antibody titres were reached already at day 3 post-boost with an immediate contraction thereafter (**Extended Data Fig. 7e**), and the overall magnitude of the increase was less pronounced (**Fig. 3d**; *P =* 0.0625). Together, these findings demonstrate that a single immunization with YF-RabG induces durable seroprotective antibody responses and establishes long-lived immunological memory that remains readily boostable months after primary vaccination.

### Dual immunity and seroprotection against yellow fever

Having established that YF-RabG induces YFV-nAbs in NHPs **(Fig. 1b; Extended Data Fig. 4b**), we next assessed the magnitude and durability of YFV-specific immunity in small-animal models. Consistent with the established immunogenicity of YF17D-based vaccines^19–21^, robust YFV-nAb responses were detected in both *Ifnar1*^−/−^ mice and Syrian golden hamsters following vaccination with YF-RabG (**Extended Data Fig. 8a,b**). In hamsters, YFV-nAb titres exhibited a clear dose-response relationship 3 weeks after immunization, with the highest vaccine dose (25x10^3^ PFU) inducing significantly higher titres than the lowest dose (10^3^ PFU; *P =* 0.0373; **Extended Data Fig. 8b**). During the one-year follow-up period, YFV-nAb titres gradually declined in all groups, with decreases below the limit of detection occurring more frequently in animals receiving lower YF-RabG doses and in comparison, with YF17D-vaccinated controls. Nevertheless, YFV-specific immunity remained detectable up to 52 weeks post-vaccination, with YFV-nAbs still measurable in 1/6, 2/6 and 4/6 animals vaccinated with 10^3^, 5x10^3^ and 25x10^3^ PFU of YF-RabG, respectively. Consistently, total YFV-specific IgG measured by indirect immunofluorescence remained detectable in 60%, 80% and 100% of animals at the respective dose levels (**Extended Data Fig. 8c**).

To determine whether YF-RabG also retained the capacity to induce YFV-specific cellular immunity, splenocytes from vaccinated BALB/c mice were stimulated *ex vivo* with the MHC class I-restricted YFV envelope peptide E_60-68_ or the MHC class II-restricted capsid peptide C_81-95_. Both YF-RabG- and YF17D-vaccinated animals exhibited robust CD8^+^- and CD4^+^-associated recall responses, reflected by increased frequencies of IFN-γ-secreting cells and elevated IFN-γ production in culture supernatants (**Extended Data Fig. 8d,e**). In contrast, no YFV-specific responses were detected in Rabipur^®^-or mock-vaccinated animals.

Together, these findings demonstrate that incorporation of the RabG transgene does not compromise the ability of the YF17D backbone to induce durable humoral and cellular immune responses against YFV. Notably, YFV-specific nAb levels can be reached and sustained that exceed those considered seroprotective in the hamster model^31^.

### Adaptation of the YF17D platform to protect against antigenically distinct lyssaviruses

Current rabies vaccines are highly effective against classical rabies virus and closely related phylogroup I lyssaviruses but generally provide limited if any protection against members of phylogroups II and III^32–34^. Amino acid sequence analysis revealed substantial divergence within key glycoprotein antigenic sites targeted by RABV-nAbs^35–37^, including antigenic sites IIb, I and III, between RABV and the phylogroup II viruses Mokola virus (MOKV) and Lagos bat virus (LBV), as well as the phylogroup III virus Lleida bat lyssavirus (LLEBV) (**Fig. 4a,b**). We therefore leveraged the adaptability of the YF17D platform to generate recombinant vaccine candidates expressing the homologous glycoproteins of MOKV, LBV or LLEBV, designated YF-MOKV-G, YF-LBV-G and YF-LLEBV-G, respectively (**Fig. 4c**).

**Fig. 4:**
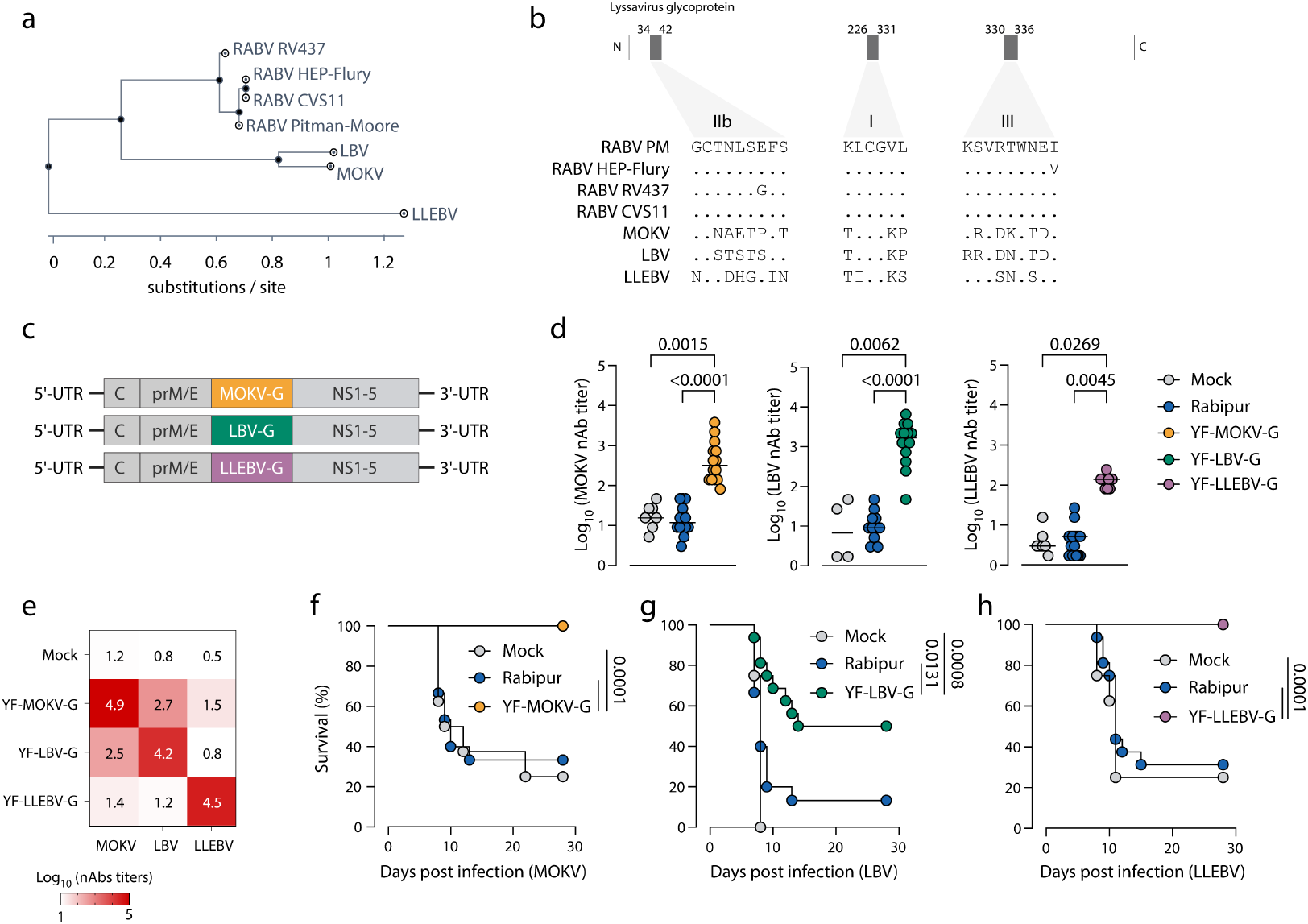
Immunogenicity and efficacy of YF-MOKV-G, YF-LBV-G and YF-LLEBV-G in mice. **a,** Glycoprotein-based phylogenetic tree of selected rabies virus (phylogroup I), Mokola virus, Lagos bat virus (phylogroup II) and Lleida bat lyssavirus (phylogroup III) sequences. Branch lengths indicate substitutions per site. **b,** Amino acid sequence alignment of the corresponding lyssavirus glycoproteins. Enlarged regions highlight substitutions within antigenic sites IIb, I and III, which contain epitopes targeted by RABV-neutralizing antibodies. **c**, Schematics of YF17D-vectored vaccine candidates encoding the glycoproteins of Mokola virus (YF-MOKV-G), Lagos bat virus (YF-LBV-G) or Lleida bat lyssavirus (YF-LLEBV-G). **d**, MOKV-, LBV-or LLEBV-neutralizing antibody titres in *Ifnar*^−/−^ mice on day 21 after vaccination (10^3^ PFU). Dots represent individual animals; bars indicate means. Dunn’s multiple-comparisons test; only P < 0.05 is shown. MOKV (Mock, n = 7; Rabipur^®^, n = 14, YF-MOKV-G, n = 14), LBV (Mock, n = 4; Rabipur^®^, n = 11, YF-LBV-G, n = 14); LLEBV (Mock, n = 7; Rabipur^®^, n = 16, YF-LLEBV-G, n = 12). **e**, Heatmap of homologous and cross-neutralizing antibody titres on day 21 after vaccination of *Ifnar*^−/−^ mice with YF-MOKV-G, YF-LBV-G or YF-LLEBV-G (10^3^ PFU, n = 6 per group). Mean titres from mock-vaccinated animals shown in panel d are included for reference. **f-h**, Survival after challenge with MOKV (f), LBV (g) or LLEBV (h). Weanling BALB/c mice (3-4 weeks old) received YF-MOKV-G, YF-LBV-G or YF-LLEBV-G (10^4^ PFU, n = 16 per group), Rabipur^®^ (1/40 human dose on days 0 and 7, n = 16 per group) or mock vaccine (2% FBS in MEM, n = 8 per group). On day 28, mice were challenged intracranially with the respective virus (50 FFU). Log-rank (Mantel–Cox) test.

All three constructs induced robust nAb responses against their respective homologous target virus in *Ifnar1*^−/−^ mice (**Fig. 4d**). In contrast, Rabipur^®^ failed to induce detectable nAb responses against MOKV, LBV or LLEBV, yielding titres comparable to those observed in mock-vaccinated animals (**Fig. 4d**).

A broader analysis of homologous and cross-neutralizing activity revealed appreciable cross-reactivity between the two phylogroup II vaccine candidates. Indeed, antibodies induced by YF-MOKV-G cross-neutralized LBV (mean log10 reciprocal titre [95% CI]: 2.8 [2.2-3.4]; P < 0.0009 *vs* mock), whereas YF-LBV-G-induced antibodies similarly neutralized MOKV (mean log10 2.6 [1.2-4.0]; *P =* 0.0114 *vs* mock) (**Fig. 4e**). In contrast, neither phylogroup II vaccine candidate elicited measurable cross-neutralization of the phylogroup III virus LLEBV, with titres remaining indistinguishable from those observed in mock-vaccinated animals (*P =* 0.1595 for YF-MOKV-G and P > 0.999 for YF-LBV-G). Likewise, sera from YF-LLEBV-G-vaccinated mice failed to significantly neutralize MOKV (mean log10 1.42 [1.3-1.6]; P > 0.999) or LBV (mean log10 1.3 [0.8-1.7]; *P =* 0.9982) relative to mock controls. Thus, while substantial serological cross-reactivity was observed within phylogroup II lyssaviruses, little evidence of cross-neutralization was found between phylogroups II and III, consistent with the greater antigenic divergence of LLEBV (**Fig. 4a,b**).

To determine whether these antibody responses translated into protection, we challenged vaccinated weanling BALB/c mice intracranially with each the corresponding homologous virus 28 days after immunization (**Fig. 4f-h**). YF-MOKV-G conferred complete protection against MOKV challenge (100% survival), whereas only 25-30% of Rabipur^®^-vaccinated and mock-vaccinated animals survived (*P =* 0.0001; **Fig. 4f**). Likewise, YF-LLEBV-G provided complete protection against LLEBV challenge (100% survival), compared with approximately 25% survival in both Rabipur^®^ and mock control groups (*P =* 0.0001; **Fig. 4h**). Protection against LBV was modest, with YF-LBV-G protecting 50% of animals following challenge, yet still significantly outperforming both Rabipur^®^ (10% survival; *P =* 0.0131) and mock- vaccinated controls (0% survival; *P =* 0.0001; **Fig. 4g**).

Together, these findings demonstrate that the YF17D platform can be readily adapted to express glycoproteins from antigenically distinct lyssaviruses, generating homologous nAb responses and conferring protection against lethal challenge beyond classical rabies virus.

## DISCUSSION

Programmes combining mass dog rabies vaccination, community engagement, surveillance and access to post-exposure care have substantially reduced dog-mediated human rabies in several endemic regions, including Latin America, the Caribbean and India^4,38–41^. However, these gains depend on sustained veterinary and healthcare infrastructure, while human prophylaxis remains constrained by repeated clinic visits, high costs and, following severe exposure, timely access to rabies vaccines and immunoglobulin (RIG)^7,15^.

The vulnerability of these control systems has become particularly evident in Europe. In the context of warfare and the complex political situation in Ukraine and neighbouring countries since 2022, disruptions to animal vaccination and surveillance, changes in domestic-animal populations and altered wildlife movement have coincided with increased rabies activity in Ukraine^42^. Concomitant with breakdown of rabies control efforts at the borders of the European Union, molecular surveillance has documented the re-emergence of sylvatic rabies variants in Moldova and several neighbouring Central European countries^43^. In parallel, quantitative modelling has identified companion-animal movement as a measurable pathway for rabies reintroduction into the European Union^44^.

These observations highlight the value of complementary human vaccines that (*i*) can establish durable immunity after fewer administrations, and (*ii*) remain compatible with licensed boosters.

Here, we show that YF-RabG, a YF17D-vectored vaccine encoding the RABV glycoprotein, combines single-dose priming with strong humoral and cellular immune responses, protection across complementary animal models, durable nAb responses and rapid recall response following administration of a licensed rabies vaccine.

As observed for our other YF17D-vectored vaccine candidates^19–21^, YF-RabG exhibited further attenuation relative to the parental YF17D virus, as indicated by lower viral titres *in vitro* and delayed mortality in suckling mice, while retaining sufficient replicative capacity to induce effective immunity. YF-RabG also retained the inserted antigen sequence for at least 11 serial *in vitro* passages, supporting its genetic stability and further development towards large-scale manufacture.

YF-RabG exhibited robust immunogenicity across multiple animal models. A single immunization elicited RABV-nAb titres above the WHO reference threshold (0.5 IU/ml) in mice, hamsters and non-human primates, whereas comparable responses required at least two administrations of Rabipur^®^. This heightened immunogenicity of YF-RabG likely derives from the intrinsic properties of the YF17D backbone. Unlike conventional inactivated rabies vaccines, which primarily induce antibody responses and CD4^+^ T-cell help^45^, YF17D is recognized for its ability to stimulate coordinated innate, humoral and cellular immunity^17,19–21^. Consistent with this biology, YF-RabG elicited robust nAb responses together with the activation of memory B-cell populations and expansion of both CD4^+^ and CD8^+^ memory T-cell compartments. Importantly, immunity directed against the YF17D backbone was preserved, indicating that insertion of RabG does not compromise the immunostimulatory properties of the vector.

The coordinated activation of B-cell and T-cell compartments may contribute to the durability and recall capacity of the response observed in this study. Although expansion of antigen-experienced B-cell populations and activation of memory T-cell subsets are consistent with the development of long-term immunological memory, the contribution of individual immune populations to protection remains to be determined. Instead, our findings support the view that YF-RabG benefits from the integrated immune stimulation characteristic of YF17D-based vaccines.

This interpretation is consistent with the durable humoral responses observed following YF-RabG vaccination. In hamsters, a single immunization maintained protective RABV-nAb levels for at least one year. Current licensed rabies vaccines, typically administered as multi-dose regimens, can induce immune memory detectable for many years, although the magnitude and persistence of the response are known to vary between individuals and can be critically influenced by factors such as immune status, age, sex, vaccine product and vaccination schedule^46–48^. Although the present animal data cannot be used to directly infer long-term protection in humans, they nevertheless demonstrate that durable rabies-specific immunity can be achieved following a single administration of YF-RabG.

Interestingly, the durability of the response differed between the rabies antigen and the YF17D vector. Whereas RABV-nAb titres remained above the accepted CoP throughout the follow-up period, YFV-nAb titres gradually declined, particularly in animals receiving lower YF-RabG doses. Despite this decline in neutralizing activity, YFV-specific binding antibodies remained detectable in most animals and were most persistent in the high-dose group. Whether such differences in nAb levels and kinetics reflect actual differences in antigen-specific immune programming with consequences for longevity of YFV-specific protective immunity and, likewise, subsequent use of YF17D-based vaccines in the presence of vector-specific antibodies remains unclear and warrants further investigation.

The establishment of durable immune memory has relevance in the context of rabies prevention. Like other prophylactic vaccines, rabies PrEP vaccination may be administered long before infection occurs, making the capacity for rapid recall a critical determinant of long-term protection. As also implemented in updated WHO guidelines^49^, properly primed immunity can alleviate from the need for an aggressive multidose PEP treatment, possibly with co-administration of RIG, after high-risk exposure. Indeed, following YF-RabG priming, administration of the licensed rabies vaccine Rabipur^®^ elicited a rapid and vigorous increase in nAb titres, demonstrating that vaccine-induced immune memory remained readily accessible for recall months after primary immunization.

From a translational perspective, this compatibility with licensed rabies vaccines may be as important as the durability of the primary response itself. Individuals receiving YF-RabG as PrEP would still be expected to receive standard-of-care post-exposure prophylaxis (PEP) following a recognized rabies exposure. Although co-administration of RIGs was not evaluated in the present study, YF-RabG was not neutralized by RABV-specific antibodies, suggesting that passive immunization may be compatible with YF-RabG even if the vaccine were to be considered for PEP.

The demonstration that YF-RabG-primed animals respond rapidly and robustly to Rabipur^®^ therefore supports integration of the platform within existing rabies prevention frameworks rather than replacement of them. If offered in time, established PEP regimens can almost completely prevent clinical rabies. Likewise, considering the almost 100% mortality of human rabies and resulting serious ethical concern that may not allow to implement any new modality for rabies PEP, YF-RabG may hence primarily be seen as a simplified priming vaccine capable of establishing protective immunity after a single administration while remaining fully compatible with currently licensed booster and PEP regimens.

This concept is supported by long-term studies of licensed rabies vaccines showing that protection depends not only on persistence of circulating antibodies but also on durable immunological memory capable of mounting rapid recall responses years after primary immunization. In a 10-year follow-up of purified chick embryo cell (PCEC)-based PrEP, 89– 98% of individuals reached the accepted protective threshold within seven days of boosting despite declining antibody concentrations over time^50^. Similarly, all participants vaccinated 10– 24 years earlier mounted an adequate response within seven days of a single booster immunization^48^. Other emerging rabies vaccine platforms point in the same direction; a single-dose adenovirus-vectored rabies vaccine recently demonstrated rapid seroconversion together with maintenance of seropositivity in most participants one year after vaccination^51^. Although the present hamster study cannot be used to predict long-term protection in humans, the rapid recall induced by Rabipur^®^ places YF-RabG within a broader framework of vaccination strategies in which durable memory responses, rather than sustained peak antibody titres alone, may support long-term protection. Such an approach could be particularly valuable in settings where completion of multi-dose vaccination schedules remains challenging.

Importantly, the ability to rapidly exchange glycoprotein antigens highlights the modularity of the YF17D platform. Current rabies vaccines provide robust protection against classical rabies virus and most phylogroup I lyssaviruses, but protection against more divergent members of phylogroups II and III remains limited^32,52^. This limitation is particularly relevant because lyssavirus diversity is maintained in wildlife reservoirs, especially bats, and includes emerging viruses such as West Caucasian bat virus (WCBV) and Lleida bat lyssavirus (LLEBV), against which current rabies biologicals offer little or no protection^52^.

The antigenic divergence observed within major neutralizing epitopes of MOKV, LBV and LLEBV was mirrored by the immunological data. Whereas strong cross-neutralization was observed between the phylogroup II viruses MOKV and LBV, little cross-reactivity was detected between phylogroups II and III. Nevertheless, replacing RabG with the corresponding glycoproteins generated immunogenic vaccine candidates that induced potent homologous nAb responses and conferred protection against homologous challenge. These findings demonstrate that the YF17D backbone can accommodate antigenically distinct lyssavirus glycoproteins without loss of immunogenicity.

Two complementary strategies are emerging for extending vaccine coverage beyond classical rabies virus. One approach is to increase breadth through antigen design, exemplified by chimeric glycoproteins such as LyssaVax, which combine domains from divergent lyssaviruses to induce broader neutralizing responses^53,54^. The other strategy is to retain a common vaccine backbone while adapting the antigen to the target virus. The present results support the latter approach and suggest that YF17D-based vectors may provide a flexible framework for responding to newly recognized lyssavirus threats. Future development may ultimately benefit from combining both concepts, pairing engineered broad-spectrum glycoproteins with live-attenuated vector platforms capable of inducing durable humoral and cellular immunity.

Collectively, our findings establish YF-RabG as a live-attenuated rabies vaccine candidate that combines single-dose priming, durable immunity and compatibility with licensed booster vaccination. We therefore envisage YF-RabG primarily as a vaccine for rabies PrEP, capable of reducing the number of administrations required to establish long-lasting protection while remaining fully compatible with existing rabies prevention strategies. At the same time, the modular nature of the YF17D platform provides a framework for the rapid development of vaccines against antigenically divergent lyssaviruses beyond classical rabies virus.

### Limitations

Several limitations should be considered when interpreting these results.

The non-human primate studies were not designed or powered to establish formal non-inferiority relative to licensed rabies vaccines. Nevertheless, the primary objective of YF-RabG is not to replace existing biologicals, but to simplify rabies PrEP through single-dose priming while preserving compatibility with current booster and PEP vaccination strategies.

Durability, measured by maintenance of nAb titres above the accepted CoP, was demonstrated for at least 70 days in non-human primates and for one year in hamsters. Although these observations cannot be directly extrapolated to humans, long-term persistence of immunity is a recognized feature of YF17D-derived vaccines. Particularly relevant prototype is the licensed live recombinant YF17D-vectored Japanese encephalitis vaccine Imojev^®^ (JE-CV), for which seropositivity remains above 90% up to a decade after a single vaccination in adults^55–57^. While YF-RabG differs from Imojev^®^ in antigen composition, construct design, and expresses an antigen derived from a different virus family as additional transgene, these clinical observations demonstrate that durable immune memory can be achieved within a YF17D-based vaccine platform and support further evaluation of YF-RabG as a long-lasting rabies priming vaccine.

The attenuation profile, apparent safety and genetic stability of YF-RabG support its further development. Notably, the YF17D platform has a long history of successful manufacturing and clinical use, including forementioned chimeric JE-VC vaccine. Nonetheless, previous studies have shown that protective efficacy can be maintained even for YF17D-derived constructs displaying some genetic instability^22^. Nevertheless, manufacturing-scale stability, reversion potential and long-term safety of YF-RabG will need to be addressed in subsequent development stages.

### Conclusion

YF-Rab is a live YF17D-vectored rabies vaccine candidate. Single-dose vaccination using YF-RabG induced potent, durable and protective immunity as demonstrated in complementary step-up models including non-human primates (immune profiling and seroprotection) and two rodent species (longevity and boostability of nAbs; protection from lethal challenge). Considering that virus-specific nAbs have been established as serological correlates of vaccine-induced protective immunity for both rabies and yellow fever, expanded non-human primate studies including lethal challenge experiments may neither be ethical nor required. Clinical evaluation will be an important next step in the development of the platform.

## MATERIALS AND METHODS

### Cells, vaccines and viruses

BHK-21J cells (baby hamster kidney fibroblasts)^58^ were kindly provided by P. Bredenbeek. BHK-21 clone 13 cells were obtained from Sigma-Aldrich (cat. no. 85011433), and Vero E6 cells (African green monkey kidney; CRL-1586) were obtained from ATCC. All cell lines were maintained in minimum essential medium (MEM; Gibco) supplemented with 10% fetal bovine serum (HyClone), 2 mM L-glutamine (Gibco) and 1% sodium bicarbonate (Gibco), and were routinely tested for mycoplasma contamination.

Licensed rabies vaccine Rabipur^®^ (GSK; lots ARBA051B, ARBA151C, ARBA551C), Ixiaro (Valneva Austria GmbH, EV20F400) and yellow fever vaccine Stamaril^®^ (Sanofi Pasteur; YF17D; lots G5400 and T3F02) were obtained through the pharmacies of University Hospitals Leuven or the Institute of Tropical Medicine Antwerp. For mouse and hamster studies, Stamaril^®^ lot G5400 was passaged three times in Vero E6 cells, and virus titres were determined by plaque assay on BHK-21J cells as described previously^59^. For the non-human primate study, Stamaril^®^ lot T3F02 was used directly without additional passaging.

YF17D virus, Stamaril^®^ (lot G5400), was passaged three times in Vero E6 cells before use. Medium used for production of virus stocks was identical to growth medium, yet containing only 2% fetal bovine serum. Virus titers were determined by plaque assay on BHK-21J cells and expressed as plaque forming units (PFU)/mL, as described before^60^.

Rabies virus isolates RV437 (GenBank accession KF154997)^61^ and CVS-11 (GenBank accession GQ918139, ATCC number: VR 959, purchased at the EURL for Rabies, ANSES, France), Mokola virus (MOKV RV610; GenBank accession HM623780.1)^62^, Lagos bat virus (LBV-D; GenBank accession JX901139)^63^ and Lleida bat lyssavirus (LLEBV, RV3208; GenBank accession NC_031955.1) ^64^ used for challenge experiments were propagated in BHK-21 cells.

### Generation of YF17D-vectored rabies and lyssavirus vaccine candidates

Vaccine constructs were generated using an infectious cDNA clone of YF17D cloned in the inducible BAC expression vector pShuttle-YF17D (patent WO2014174078A1), as previously described^59,65,66^. Briefly, the rabies vaccine candidate YF-RabG was generated by inserting the glycoprotein (G) gene of the rabies virus (RABV) Pitman-Moore vaccine strain (GenBank: AJ871962.1), corresponding to amino acids 20-524, into the full-length YF17D genome (GenBank: X03700) as an in-frame translational fusion within the E/NS1 intergenic region^18,67^. The RabG sequence was fused to an additional transmembrane domain derived from West Nile virus (WNV, GenBank accession AF196835.2) envelope protein to preserve the topology of the YF17D polyprotein and enable proper endoplasmic reticulum luminal expression of the heterologous antigen^20,68^ (**Extended Data Fig. 1a**).

Recombinant viruses were rescued by transfection of BHK-21 cells with the corresponding BAC constructs. Virus-containing supernatants were harvested 4 days post-transfection upon the appearance of extensive cytopathic effect and titrated by plaque assay on BHK-21J cells using an agarose overlay, as previously described^59,60^. Viral titers were expressed as plaque-forming units (PFU)/ml. The integrity of the inserted transgene was confirmed by Sanger sequencing of rescued virus stocks.

Additional YF17D-based vaccine candidates expressing glycoproteins from Mokola virus (MOKV), Lagos bat virus (LBV) and Lleida bat lyssavirus (LLEBV), designated YF-MOKV-G, YF-LBV-G and YF-LLEBV-G, respectively, were generated using the same cloning strategy. The MOKV glycoprotein gene (GenBank: HM623780.1) was amplified by PCR from the pI.18-MOKV-G expression plasmid^69^, whereas LBV (GenBank: GU170202.1) and LLEBV (GenBank: NC_031955.1) glycoprotein genes were amplified by RT-PCR from viral RNA. The resulting glycoprotein sequences were inserted into the YF17D E/NS1 intergenic region as described above for YF-RabG.

### Virus growth kinetics

Virus growth kinetics were assessed in BHK-21J cells infected at a multiplicity of infection (MOI) of 0.01. After 1 h incubation at room temperature (RT), inocula were removed, cells were washed, and fresh medium was added. Culture supernatants were collected daily for 4 days and stored at -80 °C until analysis. Viral titers were determined by endpoint titration on BHK-21J cells and calculated using the Reed-Muench method with the Lindenbach calculator^70^. Titers were expressed as 50% tissue culture infectious doses (TCID_50_/ ml).

### Immunofluorescence analysis of antigen expression

Antigen expression by YF-RabG-infected cells was assessed by immunofluorescence staining as previously described^21,59^. Briefly, BHK-21J cells were infected with YF-RabG and fixed at 2 days post-infection with 4% paraformaldehyde for 20 min. For intracellular antigen detection, cells were permeabilized with 0.1% Triton X-100 for 5 min. Following washing with PBS, cells were blocked with 1% fetal bovine serum (FBS) in PBS containing 0.05% Tween-20 for 1 h at RT.

Yellow fever virus antigens were detected using polyclonal hamster anti-YF17D serum derived from Syrian hamsters vaccinated with YF17D (Stamaril^®^; 1:100). Rabies virus glycoprotein (RabG) expression was detected using a mouse monoclonal anti-rabies glycoprotein antibody (MyBioSource, MBS310434, 1:100, or Thermo Fisher Scientific, MA1-7079, 1:100). To visualize surface-expressed RabG, staining was additionally performed on non-permeabilized cells. Following incubation with primary antibodies, cells were washed and incubated with goat anti-mouse Alexa Fluor 594 (Life Technologies; 1:500) and rabbit anti-hamster FITC (Jackson ImmunoResearch; 1:500) secondary antibodies. Nuclei were counterstained with DAPI (Sigma-Aldrich).

All confocal images were acquired using identical imaging settings on a Leica TCS SP5 confocal microscope equipped with an HCX PL APO 63×/1.2 NA water-immersion objective.

### Genetic stability analysis

To assess the genetic stability of YF-RabG during serial propagation in cell culture, virus recovered from transfected BHK-21 cells (P0) was passaged sequentially up to passage 11 (P11). At each passage, fresh BHK-21 cells were inoculated for 1 h at RT with a 1:10 dilution of supernatant from the preceding passage. Cells were subsequently washed three times with culture medium and incubated for 48-72 h before supernatants were harvested for the next passage.

Retention of the inserted transgene was assessed at each passage by RNA extraction and DNase I treatment (Direct-zol RNA Miniprep Kit, Zymo Research), followed by RT–PCR (qScript XLT One-Step RT–PCR Kit, Quanta Biosciences) and Sanger sequencing.

### Plaque reduction neutralization assay (PRNT)

PRNTs were performed using immune sera directed against either RABV or YFV. YF-RabG was incubated for 1 h at 37 °C with serial dilutions of archived serum from Rabipur^®^-immunized mice (RABV-neutralizing antibody titre of 17 IU/ml) or archived serum from Stamaril^®^-immunized non-human primates. Normal mouse and non-human primate sera were included as negative controls. Following incubation, serum-virus mixtures were added to confluent BHK-21J cell monolayers seeded in 6-well plates and allowed to adsorb for 1 h at RT. Cells were subsequently washed twice with PBS and overlaid with MEM containing 0.5% ultrapure low-melting-point agarose (Invitrogen). Plates were incubated at 37 °C for 6 days, after which cells were fixed with 4% formaldehyde and plaques were visualized by staining with 1% crystal violet. Plaques were counted manually and expressed as the number of plaques per well.

### Animals

All procedures involving experimental animals were approved by the relevant institutional animal ethics committees, conducted in accordance with applicable national regulations and FELASA guidelines, and reported herein in accordance with ARRIVE (Animal Research: Reporting of In Vivo Experiments) guidelines. Sample sizes were defined *a priori* by performing a power analysis with the expected effect size, statistical power of 80% and a significance level of 0.05, using G*Power^71^.

Unless otherwise specified, animals were inspected at least twice daily throughout the study. Prior to experimental procedures, assessments focused on general health and welfare, including body condition, posture, grooming behaviour and activity. During experimental procedures, animals were monitored daily for the development of clinical signs, adverse events or disease-related symptoms. Experiment-specific humane endpoints and clinical scoring criteria are described in the section *Animal experiments*. Animals reaching predefined humane endpoints were euthanized in accordance with institutional and ethical guidelines.

#### Mice

Three-to five-day-old BALB/c pups used for neurovirulence studies were obtained together with foster dams from Janvier Laboratories (Le Genest-Saint-Isle, France). Specific-pathogen-free (SPF) BALB/c mice used for immunization and challenge studies were purchased from Janvier Laboratories (6–8 weeks old) or Charles River Laboratories (Margate, UK; 3–4 weeks old, female mice). Type I interferon receptor-deficient mice (*Ifnar1*^−/−^, B6.129S2-*Ifnar1^tm1Agt^*)^29^ were bred in-house at the Rega Institute, KU Leuven.

Mice were housed in individually ventilated (IsoCage N, Tecniplast) or filter-top cages, with a maximum of five per cage, under specific pathogen-free conditions with a 12-h light–dark cycle (room temperature: 22 °C ± 2, humidity: 45–70%). Food and water were provided *ad libitum* and environmental enrichment, including nesting material, cotton, and cardboard tunnels or shelters. Bedding material was weekly renewed and all materials were autoclaved. Purchased mice were allowed to acclimatise for one week prior to starting the experiment. Formal randomisation was not used for treatment allocation. To minimise potential bias, cage positions on the rack were not assigned according to treatment group, cages were coded, and personnel responsible for clinical assessments were blinded to treatment allocation throughout the study.

Studies were approved by the institutional ethical committees of KU Leuven (licenses P170-2017, P100-2019 and P115-2021) and Sciensano (licenses 20161103, 20171207 and 20210301). Experiments conducted at the Animal and Plant Health Agency (APHA) were performed in ACDP3/SAPO4 containment facilities in accordance with the UK Animals (Scientific Procedures) Act 1986 under Home Office licence PCA17EA73.

#### Hamsters

Female Syrian golden hamsters (*Mesocricetus auratus*; RjHan:AURA, 6–9 weeks old) were obtained specific pathogen-free from Janvier Laboratories (Le Genest-Saint-Isle, France) and allowed to acclimatise for one week prior to starting the experiment. Animals were housed in individually ventilated isolator cages (maximum two per cage) with food and water provided *ad libitum* and environmental enrichment (wood blocks and cotton nesting material). Bedding material was weekly renewed and all materials were autoclaved.

All hamster studies were conducted at the Rega Institute, KU Leuven. Experimental procedures and housing conditions were approved by the KU Leuven Ethical Committee (licence P170-2017).

#### Non-human primates

Twelve outbred adult female cynomolgus macaques (*Macaca fascicularis*) were included in this study. Animals were purpose-bred and housed at the Biomedical Primate Research Centre (BPRC; Rijswijk, the Netherlands). All animals were deemed healthy based on physical examination and baseline clinical chemistry and haematological parameters.

All procedures and animal husbandry were conducted at BPRC following approval by the Dutch Central Committee for Animal Experiments (CCD, project licence AVD5020020171326) and by the institution’s Animal Welfare Body.

### Animal experiments

#### Vaccine virus safety assessment

BALB/c suckling mice (3–5 days old; n = 11 per group) were inoculated intracranially with 10 μl of virus suspension containing either 10 PFU of YF17D or 1,000 PFU of YF-RabG. Control animals received culture medium alone (mock). Inoculations were performed using a Micro-Fine™ insulin needle (BD Biosciences). Following inoculation, animals were monitored daily for 11 days for survival, mobility and the development of neurological signs. Humane endpoints included severe neurological impairment, inability to move normally, failure to nurse, or moribund condition, at which point animals were euthanized by intraperitoneal administration of sodium pentobarbital (Dolethal, 200 mg/ml; Vétoquinol SA).

#### BALB/c immunization and challenge studies

To assess vaccine immunogenicity, BALB/c mice (6–8 weeks old) were immunized with a single dose of YF-RabG (2x10^3^ PFU, i.p.). To permit replication of YF17D-based vaccine viruses in mice, animals receiving YF-RabG or YF17D were administered anti-IFNAR antibody (MAR1-5A3; 2 mg per mouse in 100 µl PBS, i.p.) 4 h before vaccination. Control groups received either YF17D (2x10^3^ PFU, i.p.), Rabipur^®^ (25 µl, equivalent to 1/40 human dose, administered i.p. on days 0 and 7), or MEM supplemented with 2% FBS (mock control, i.p.). Blood samples were collected under isoflurane anaesthesia from the submandibular vein 28 days post-vaccination for determination of rabies virus (RABV)-neutralizing antibody titres. Mice were subsequently euthanized by intraperitoneal administration of sodium pentobarbital (Dolethal, 200 mg/ml) and spleens harvested for cellular immune analyses.

To assess short- and longer-term protection, separate cohorts of vaccinated animals were challenged either 28 or 90 days after immunization. Animals were inoculated intramuscularly (i.m.) with RABV isolate RV437 (720 TCID_50_ in 80 μl; 40 μl per hind limb). Animals were monitored daily for 5 weeks for survival, body weight and the development of clinical signs of rabies. Humane endpoints included marked deterioration of body condition, persistent inactivity, neurological abnormalities such as impaired or circular gait, hind limb paresis/paralysis, inability to access food or water, or moribund condition. Animals reaching predefined humane endpoints were euthanized by intraperitoneal administration of sodium pentobarbital (Dolethal, 200 mg/ml; Vétoquinol SA). Clinical scoring was based on body weight, body condition, behaviour and physical appearance.

To evaluate protection against phylogroup II and divergent lyssaviruses, female BALB/c mice (3–4 weeks old) were vaccinated with a single dose (10^4^ PFU, i.p.) of YF-MOKV-G, YF-LBV-G or YF-LLEBV-G. Animals received anti-IFNAR antibody (MAR1-5A3; 2 mg per mouse, i.p.) 4 h before vaccination. Control groups received either MEM supplemented with 2% FBS (mock control) or a single dose of Rabipur^®^ (1/40 human dose, i.p.). Blood samples were collected 21 days post-vaccination for determination of virus-neutralizing antibody titres. At day 28, animals were challenged intracranially with the corresponding lyssavirus (MOKV, LBV, LLEBV, 50 FFU in 30 µl) and monitored for 4 weeks.

In a separate efficacy study, BALB/c mice (3–4 weeks old) received Rabipur^®^ (1/40 human dose, i.p.) or a single dose of YF-RabG (10^4^ PFU, i.p.) following anti-IFNAR treatment (MAR1-5A3; 2 mg per mouse, 4 h before vaccination, i.p.). Mock-vaccinated animals received MEM supplemented with 2% FBS. At day 28 post-vaccination, mice were challenged intracranially with RABV strain CVS-11 (50 FFU in 30 μl) and monitored for 4 weeks.

Prior to intracranial challenge, animals were anaesthetized with isoflurane/oxygen and received buprenorphine (0.05 mg/kg, s.c.) for periprocedural analgesia. Humane endpoints were defined as affected gait (slow or circular movements) or hind quarter paralysis. Animals reaching humane endpoints were euthanized by exsanguination under terminal isoflurane/oxygen anaesthesia, and blood was collected by cardiac puncture^72^.

#### Ifnar1^−/−^ immunization and challenge studies

*Ifnar1^−/−^* mice (6–8 weeks old) were immunized with a single dose of YF-RabG (10^3^ PFU, i.p.). Control groups received either Rabipur^®^ (1/40 human dose administered on days 0 and 7 together with Ixiaro^®^ on day 0, i.p.) or Ixiaro^®^ alone. Ixiaro^®^ was included as an unrelated inactivated flavivirus vaccine control. Blood samples were collected under isoflurane anaesthesia from the submandibular vein on days 14, 21 and 28 post-vaccination for determination of RABV-neutralizing antibody titres.

At 2 months post-vaccination, mice were challenged intramuscularly with RABV isolate RV437 (720 TCID_50_ in 80 μl; 40 μl per hind limb). Animals were monitored daily for 5 weeks and euthanized upon reaching predefined humane endpoints, as described above for the short- and longer-term protection experiments.

To evaluate the immunogenicity of YF17D-vectored lyssavirus vaccines, *Ifnar1*^−/−^ mice (6–8 weeks old) were immunized with a single dose (10^3^ PFU, i.p.) of YF-MOKV-G, YF-LBV-G or YF-LLEBV-G. Control groups received either Rabipur^®^ (1/40 human dose, i.p.) or MEM supplemented with 2% FBS (mock control). Blood samples were collected under isoflurane anaesthesia from the submandibular vein 21 days post-vaccination for determination of virus-neutralizing antibody titres.

#### Hamster immunization studies

Longevity study: Hamsters were immunized with a single dose of YF-RabG (10^3^, 5x10^3^ or 25x10^3^ PFU; i.p.). A control group received YF17D (5x10^3^ PFU; i.p.). Blood samples were collected via the jugular vein under isoflurane anaesthesia at day 21 and at 6 and 12 months post-vaccination for determination of RABV- and YFV-neutralizing antibody titres. YFV-specific binding antibodies at 6 and 12 months post-vaccination were assessed by indirect immunofluorescence assay using commercial slides (EUROIMMUN; FI 2665-1005 G, Lübeck, Germany).

Boostability study: Hamsters were immunized with a single dose of YF-RabG (5x10^3^ PFU; i.p.) or a YF17D (5x10^3^ PFU; i.p.). A control group received Rabipur^®^ (1/40 human dose; days 0 and 7, i.p.). Blood samples were collected via the jugular vein under isoflurane anaesthesia 21 days post-vaccination for determination of RABV-neutralizing antibody titres.

Sixteen weeks after primary immunization, YF-RabG- and Rabipur^®^-primed animals received a booster dose of Rabipur^®^ (1/40 human dose, i.p.). Serum samples were collected immediately before boosting (day 0) and at days 1, 3, 7 and 14 thereafter for determination of RABV-neutralizing antibody titres. Age-matched naïve hamsters immunized with a single dose of Rabipur^®^ (1/40 human dose, i.p.) were included as controls for primary seroconversion kinetics and sampled at the same time points.

#### Immunization of non-human primates

Twelve healthy adult female cynomolgus macaques (*Macaca fascicularis*; ≥4 years of age; ≥4.1 kg body weight) were screened before study initiation and confirmed seronegative for YFV, related flaviviruses and RABV. Animals were pair-housed with socially compatible cage mates and randomly assigned to two groups (n = 6 per group).

Macaques were immunized with YF-RabG (10^5^ PFU; s.c.) on day 0 and received a homologous booster dose on day 28. Control animals received Stamaril^®^ (one human dose; s.c., day 0) and Rabipur^®^ (one human dose; i.m., days -7 and 0), followed by booster immunizations with the respective vaccine(s) on day 28.

Animals were monitored daily for health status, appetite, general behaviour and stool consistency. Blood samples were collected longitudinally for haematological and clinical chemistry analyses, as well as for assessment of humoral and cellular immune responses. Follow-up was continued until day 70 post-vaccination.

### Rabies virus rapid fluorescent focus inhibition test (RFFIT)

RABV-neutralizing antibody titres were determined using the rapid fluorescent focus inhibition test (RFFIT) according to the World Organisation for Animal Health (WOAH; formerly OIE) guidelines. Briefly, heat-inactivated serum samples were serially diluted and incubated with a predetermined dose of rabies virus strain CVS-11. BHK-21 cells were subsequently added to the serum-virus mixtures and plates were incubated for 22 ± 1 h at 37 °C in 5% CO_2_. Cells were then fixed and stained with a fluorescein isothiocyanate (FITC)-conjugated anti-rabies antibody (FDI-Fujirebio Diagnostics) to detect non-neutralized virus. Infected fields were scored by fluorescence microscopy and neutralizing antibody titres were calculated using the Reed-Muench method with the Lindenbach calculator^70^. Titres were expressed as IU/ml relative to the Second International Standard for Rabies Immunoglobulin (European Union Reference Laboratory for Rabies, ANSES, Nancy, France).

### Modified fluorescent antibody virus neutralization test (mFAVN) for non-RABV lyssaviruses

Neutralizing antibody titres against non-RABV lyssaviruses were determined using a modified fluorescent antibody virus neutralization test (mFAVN)^73,74^. In brief, serum samples were heat-inactivated (56 °C for 30 min), serially diluted and incubated with 100 TCID_50_ of the respective lyssavirus before addition of BHK-21 cells. Following incubation, infected cells were detected by fluorescent anti-rabies antibody staining and neutralization titres were calculated as IC_50_ values.

### YFV neutralization test

To quantify YFV-specific neutralizing antibodies, serum samples were tested in a validated fluorescence-based seroneutralization assay (SNT) using an mCherry-tagged YF17D reporter virus as previously described^75^. In brief, serum samples were heat-inactivated (56 °C for 30 min), serially diluted and incubated with the reporter virus for 1 h at 37 °C before transfer onto BHK-21J. The virus input was predetermined by back-titration to yield 1,500–2,000 spot-forming units per well (MOI = 0.02). Cells were incubated for 3 days at 37 °C, after which cells were fixed with 4% formaldehyde. Fluorescent spots were scanned and counted using a CTL ImmunoSpot S6 Ultimate reader (FluoroSpot/BioSpot modes). Dose-response curves were normalized to these controls and fitted in Genedata Screener (Genedata AG, version 17.0.4) to determine the serum dilution reducing spot counts by 50% (SNT_50_).

### IFN-γ recall responses in mouse splenocytes (ELISpot and ELISA)

YF17D- and RabG-specific cellular immune responses were evaluated 28 days post-vaccination by IFN-γ ELISpot and ELISA. Splenocytes were isolated by mechanical dissociation through a 70-μm cell strainer followed by red blood cell lysis (eBioscience). For ELISpot analysis, 5x10^5^ splenocytes per well were seeded onto mouse IFN-γ FluoroSpot plates (Mabtech) and stimulated for 16 h with individual synthetic peptides corresponding to YFV MHC class I (E_60−68_, CYNAVLTHV) and MHC class II (C_81−95_, RKVKRVVASLMRGLS)^76^ epitopes, or RabG MHC class I (RabG_453−461_, LPNWGKYVL) and MHC class II (RabG_311−330_, TTKSVSFRRLSHLRKLVPGF) epitopes. Medium supplemented with DMSO served as a negative control. Plates were developed according to the manufacturer’s instructions and spot-forming units (SFU) were quantified using an ImmunoSpot S6 Universal reader (CTL). Data are expressed as SFU after subtraction of background values obtained from unstimulated control wells.

For IFN-γ quantification by ELISA, splenocytes were stimulated under the same conditions and culture supernatants were collected after incubation. IFN-γ concentrations were determined using a commercial mouse IFN-γ ELISA kit (Mouse IFN gamma ELISA Ready-SET-Go! Kit, BMS606TWO, Thermo Fisher Scientific) according to the manufacturer’s instructions. Background values obtained from unstimulated control wells were subtracted from stimulated samples, and negative values were set to zero.

### Flow cytometric analysis of PBMCs in non-human primates

Peripheral blood (200 μl) collected in EDTA was stained for 20 min at RT with an antibody panel including markers for B-cell identification (CD20, CD21, CD27 and IgD), T-cell characterization (CD3, CD4, CD8, CD28, CD95, CD25 and CD127), innate immune populations (CD14, CD16, CD11c, CD1c, CD123 and CD163), and cellular activation (HLA-DR, CD40, CD69, CD71, CD80, CD83 and CD86). Fluorophores, antibody clones and working dilutions are provided in **Supplementary Table 1**. Erythrocytes were lysed using BD Pharm Lyse™ (BD Biosciences) for 10 min, after which cells were washed, fixed in 2% paraformaldehyde and acquired on a Cytek Aurora spectral flow cytometer (Cytek Biosciences, Fremont, CA, USA). Data were analysed using FlowJo v10.8.2 (BD Biosciences).

### High-dimensional analysis of flow cytometry data

For high-dimensional analyses, CD45^+^ cells were manually gated from individual samples and 10^4^ randomly selected cells per sample were concatenated into a single dataset. t-distributed stochastic neighbour embedding (t-SNE) maps were generated using canonical lineage markers (CD3, CD20, CD4, CD8, HLA-DR, CD14, CD16, CD11c and CD1c) with the opt-SNE algorithm77 and the following parameters: 1,000 iterations, perplexity = 3, KNN = exact and gradient = Barnes-Hut.

Unsupervised clustering was performed using FlowSOM^78^ with the same marker set, using a 10 × 10 self-organizing map and 10 meta-clusters. FlowSOM heatmaps were used to assign cell identities and define major immune cell populations (**Extended Data Fig. 5**). FlowSOM clusters were subsequently grouped into B cells, CD4^+^ T cells, CD8^+^ T cells, dendritic cells and monocytes for downstream analyses.

For subset-specific analyses, t-SNE mapping was repeated on individual B cell and T cell compartments using only lineage, phenotypic and activation markers relevant to the respective cell population.

### Statistical analysis

Statistical analyses were performed using GraphPad Prism 10 software (GraphPad Software). Sample sizes, statistical tests and measures of central tendency are indicated in the corresponding figure legends.

Comparisons between two groups were performed using two-sided Mann-Whitney tests or two-tailed paired t-tests, as indicated. Comparisons involving more than two groups were performed using Kruskal-Wallis tests followed by Dunn’s multiple-comparison test or two-way ANOVA followed by Tukey’s multiple-comparison test, as appropriate. Longitudinal datasets were analyzed using a mixed-effects model fitted by restricted maximum likelihood (REML) with Geisser-Greenhouse correction, followed by Šídák’s multiple-comparisons test. Survival curves were compared using the log-rank (Mantel-Cox) test.

For ELISpot and ELISA analyses, background values obtained from unstimulated control samples were subtracted from stimulated samples and negative values were set to zero. All tests were two-sided and P < 0.05 was considered statistically significant.

## ACKNOWLEDGEMENTS

We thank Jasmine Paulissen, Dr. Winnie Kerstens, Sofie Leen, Hooman Goharriz and David Seldon for their contributions to the serological assessments. Dr. Sapna Sharma, Carolien De Keyzer, Lindsey Bervoets, Aurélie Francart, Leigh Thorne and David Seldon supported *in vivo* studies; Charlotte Vanderheydt supported project management. We also acknowledge the staff of the animal facilities at BPRC, APHA and Sciensano and the KU Leuven Rega Institute for their excellent scientific-technical support. We are grateful to Dr. Charles Rupprecht, Dr. Dirk E. Teuwen and Dr. Maarten Leyssen as members of the Scientific Advisory Board of RABYD-VAX for the fruitful scientific discussions and critical guidance throughout the project.

## AUTHOR CONTRIBUTIONS (CRediT)

**Conceptualization:** KD, LC, JN, PL, ACB, ARF, SVG, CV

**Data curation:** LSF, YAA, KG

**Formal analysis:** LSF, YAA, ST

**Funding acquisition:** LC, PL, JN, KD, ARF, ACB, OD, SVG, EV, MR

**Investigation:** LSF, ST, GW, BEV, HGT, PPL, GK, KPB, NM, SD, KG, JM

**Methodology:** LSF, YAA, ST, GW, BEV, HGT, PPL, GK, KPB, NM, SD, KG, JM, TV, HJT, MR

**Project administration:** LC, PL

**Resources:** KD, JN, ARF, OD, SVG, MR, EV, WB

**Supervision:** KD, LC

**Visualization:** LSF, YAA

**Writing –original draft:** LSF, YAA, LC, KD

**Writing –review & editing:** all authors

## DATA AVAILABILITY

Datasets generated and/or analyzed during the current study can be provided upon reasonable request to the corresponding authors.

