## Supplementary Material for "Seroprotection and improved polyfunctional immunity conferred by a YF17D-vectored rabies vaccine"

**List of Supplementary materials**

Extended Data Figures 1-8

Supplementary Table 1

EXTENDED DATA FIGURE 1

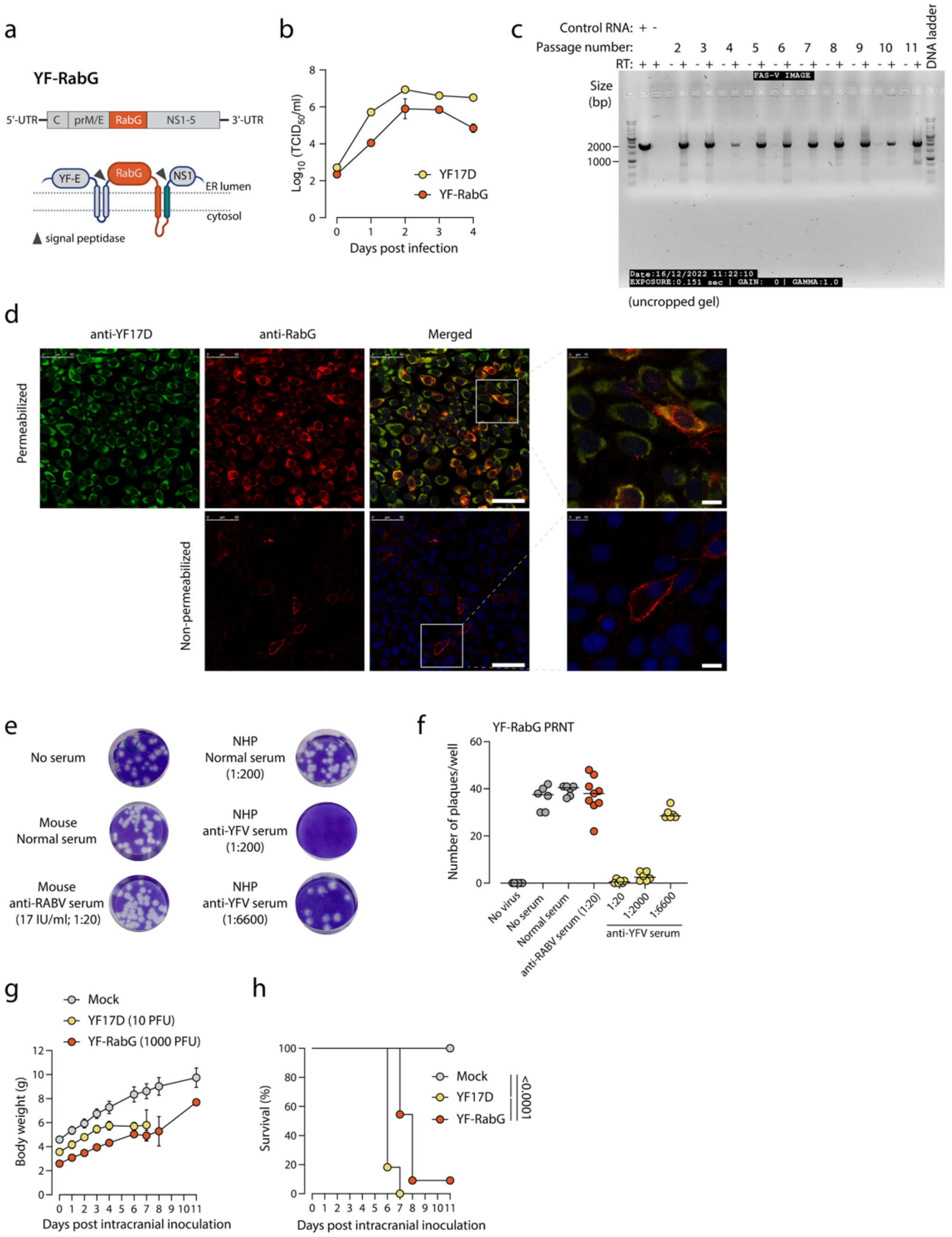


**Extended Data Fig. 1: Vaccine design and *in vitro* characterization of the YF17D-vectored rabies vaccine candidate YF-RabG.**

**a,** Schematic of YF-RabG. The rabies virus glycoprotein (RabG; orange), lacking the N-terminal signal peptide, was inserted into the E/NS1 intergenic region as a translational fusion within the YF17D polyprotein. To maintain proper folding topology, an additional transmembrane domain derived from West Nile virus E protein (green) was fused to the C terminus of RabG.

**b**, Growth kinetics of YF17D and YF-RabG in BHK-21 cells infected at a multiplicity of infection (MOI) of 0.01. Viral titers were determined over time by endpoint titration on BHK-21 cells. Data are mean ± s.e.m. (YF17D, n = 7; YF-RabG, n = 5).

**c**, RT-PCR analysis of viral RNA from YF-RabG serial passages (P2-P11), performed in the presence (+RT) or absence (–RT) of reverse transcriptase. Parental YF-RabG cDNA served as a positive control and $ddH_2O$ as a negative control. A 1-kb DNA ladder was used as a molecular weight marker.

**d**, Antigenicity of YF-RabG. Confocal immunofluorescence of BHK-21 cells at 2 days post-infection with YF-RabG. Cells were stained for YF17D (green) and RabG (red) after permeabilization (top), or RabG only without permeabilization. Nuclei were counterstained with DAPI (blue). Scale bars: 50 µm; insets, 10 µm.
**e**, Representative plaque reduction neutralization test (PRNT) in cells infected with YF-RabG. Prior to infection, the virus was preincubated with preimmune serum (Normal serum), or immune sera from animals vaccinated with Rabipur® (anti-RABV serum) or Stamaril® (anti-YFV serum) at the indicated dilution.
**f**, Number of plaques per well. The horizontal bars represent the median values.
**g**, **h**, Weight change (c) and survival (d) of suckling BALB/c mice (3-5 days old) following intracranial inoculation with medium (Mock, n = 11), 10 PFU YF17D (n = 11), or 1000 PFU YF-RabG (n = 11). Survival was analyzed by log-rank (Mantel–Cox) test.

#### EXTENDED DATA FIGURE 2

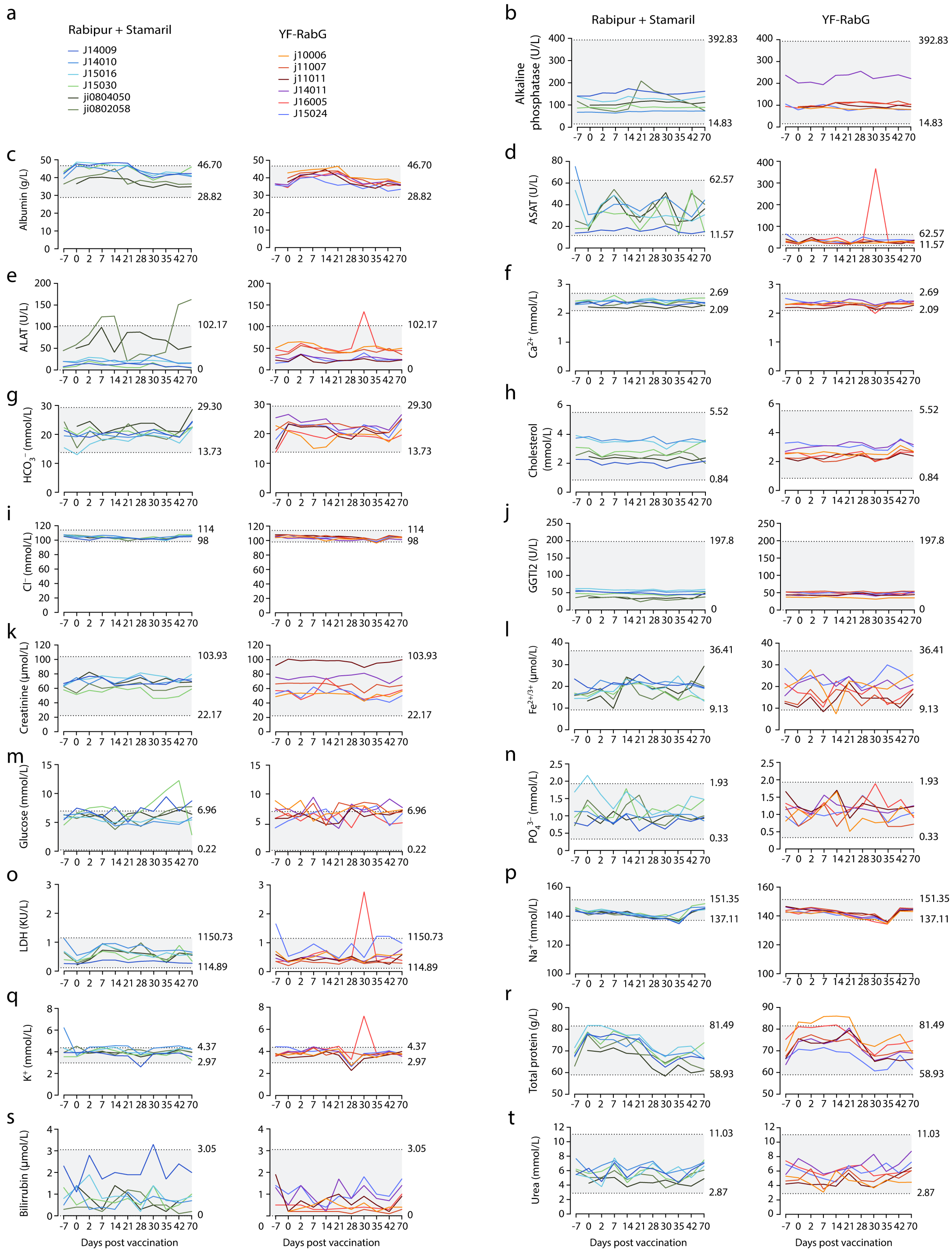


**Extended Data Fig. 2: Clinical chemistry monitoring of immunized macaques.**

**a**, Animal identifiers for each experimental group.

**b–t**, Blood biochemical parameters measured over time following vaccination. Concentrations of alkaline phosphatase, albumin, aspartate aminotransferase (ASAT), alanine aminotransferase (ALAT), calcium, bicarbonate, cholesterol, chloride, gamma-glutamyl transferase (GGTI2), creatinine, iron, glucose, phosphate, lactate dehydrogenase (LDH), sodium, potassium, total protein, total bilirubin and urea are shown for individual animals. Shaded areas indicate reference ranges for healthy individuals.

#### EXTENDED DATA FIGURE 3

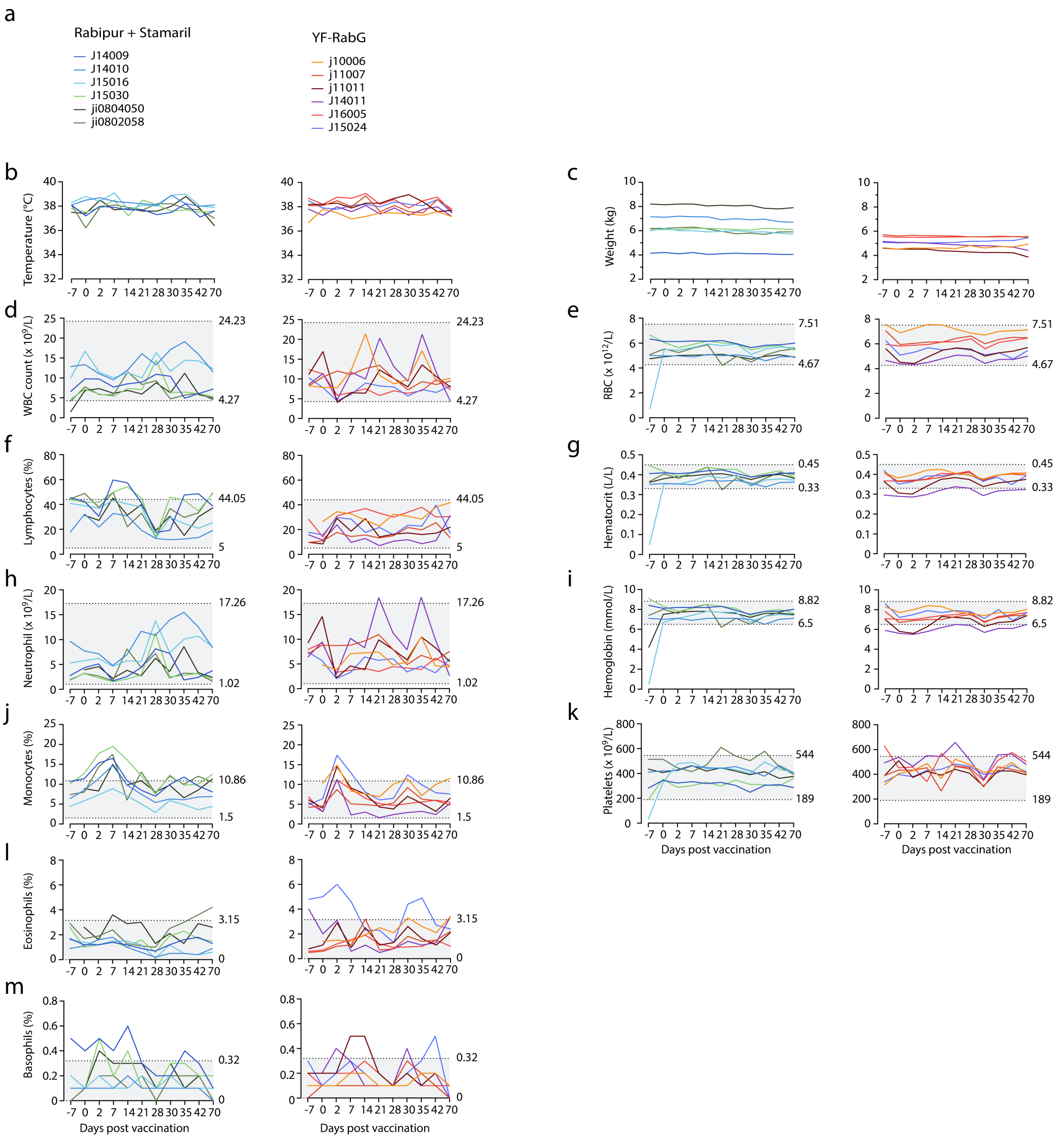


**Extended Data Fig. 3: Clinical and hematological monitoring of immunized macaques.**
**a**, Animal identifiers for each experimental group.
**b,c**, Body temperature (b) and weight (c) measured over time following vaccination.
**d–m**, Hematological parameters measured longitudinally. White blood cell (WBC) and neutrophil counts, percentages of lymphocytes, monocytes, eosinophils and basophils, red blood cell (RBC) count and platelet count, hematocrit and hemoglobin levels are shown for individual animals. Shaded areas indicate reference ranges for healthy individuals.

#### EXTENDED DATA FIGURE 4

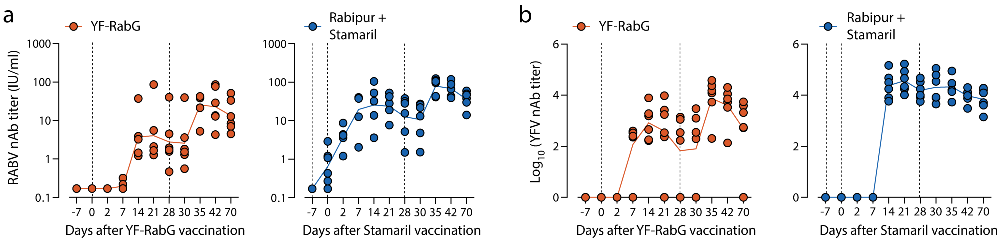


**Extended Data Fig. 4:** Kinetics of the neutralizing antibody response in NHPs after vaccination.
**a, b,** Neutralizing antibody titers against rabies virus (RABV, a) and yellow fever virus (YFV, b) at indicated time points post-vaccination. Data points represent individual animals, and lines indicate geometric mean titers (GMTs). Vertical dashed lines denote vaccination time points.

**EXTENDED DATA FIGURE 5**

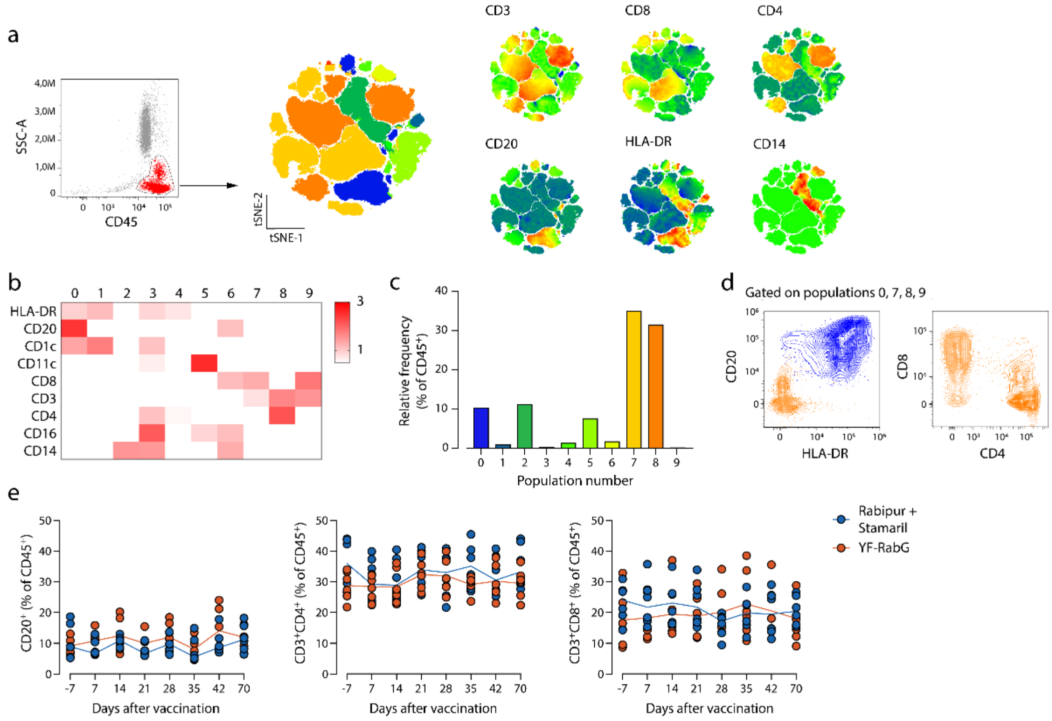


**Extended Data Fig. 5**: **Unbiased cell clustering strategy for flow cytometry data analysis of PBMCs.**

**a,** $CD45^+$ cells were manually gated from individual samples, concatenated, and analyzed using opt-SNE based on canonical surface markers for B cells, T cells, monocytes and antigen-presenting cells (APCs). Cell populations were defined by unsupervised clustering using FlowSOM. Relative mean fluorescence intensity of CD3, CD8, CD4, CD20, HLA-DR and CD14 is shown across the t-SNE space.

**b,** Relative expression of phenotypic surface markers across FlowSOM-defined populations. Values are Z-scored within each marker.

**c,** Proportion of $CD45^+$ cells assigned to each FlowSOM population.

**d,** Contour plots of selected populations (0, blue; 7, light orange; 8, dark orange) using canonical B cell (left) and T cell (right) markers.

**e,** Frequencies of circulating $CD20^+$, $CD4^+$ T and $CD8^+$ T cells among $CD45^+$ cells. Lines indicate mean values; points represent individual animals.

### EXTENDED DATA FIGURE 6

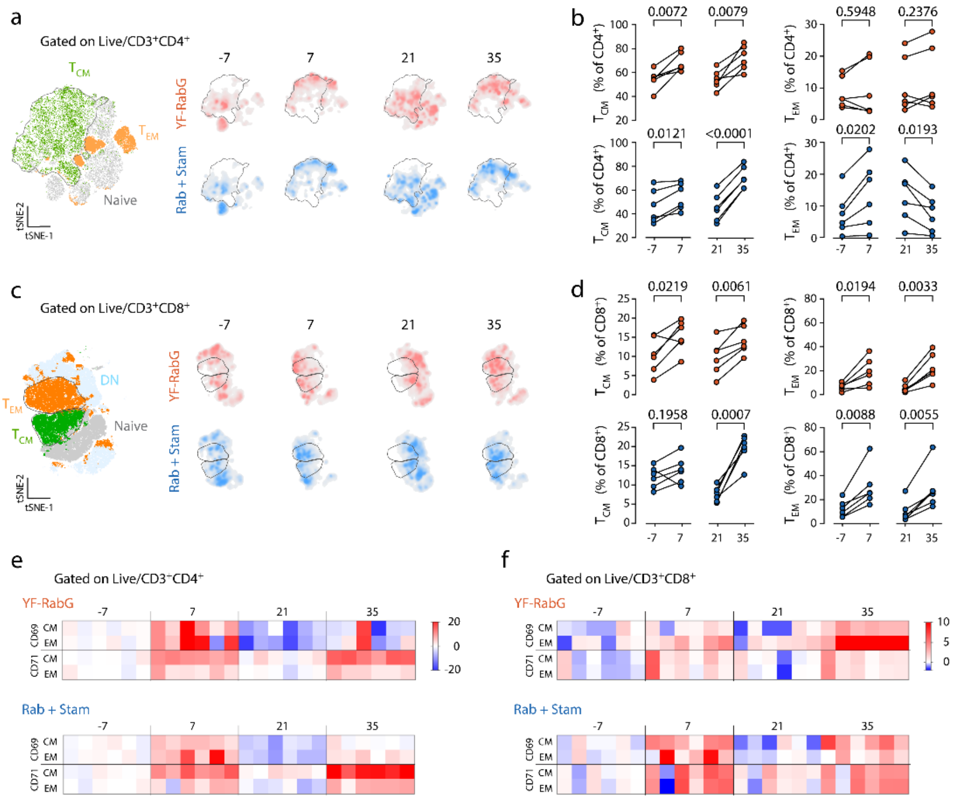


**Extended Data Fig. 6: Characterization of circulating memory T cells in NHPs after YF-RabG vaccination.**

**a, c,** t-SNE maps of CD4$^{+}$ (a) and CD8$^{+}$ (c) T cells showing clustering of central and effector memory subsets and their density distribution in t-SNE space across time points.

**b, d,** Frequencies of circulating central and effector memory CD4$^{+}$ (b) and CD8$^{+}$ (d) T cells. Points represent individual animals. P values were calculated using two-tailed paired t-tests.

**g,** Heatmap of activation marker expression (CD69, CD71), z-scored to day −7.

#### EXTENDED DATA FIGURE 7

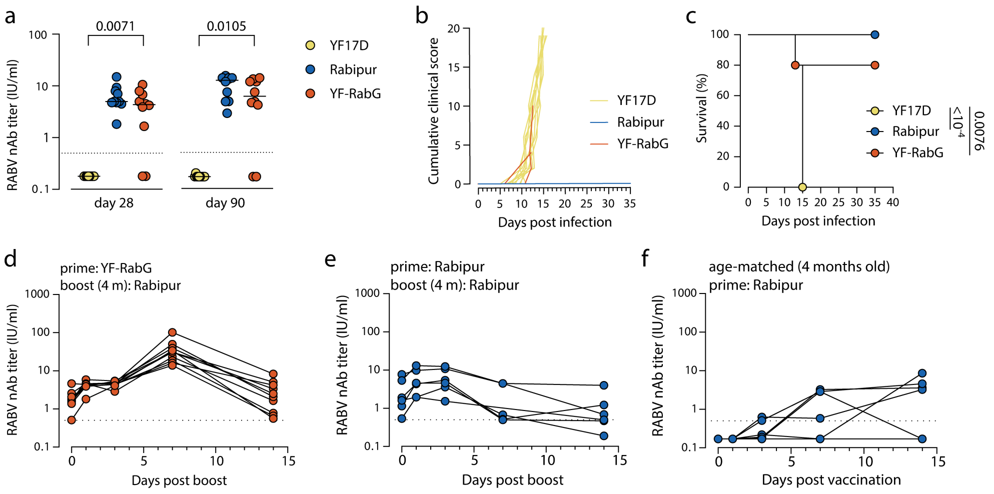


**Extended Data Fig. 7: Neutralizing antibody responses, protective efficacy and boostability of YF-RabG.**

**a,** RABV-neutralizing antibody titers in adult BALB/c mice measured by RFFIT at days 28 and 90 post-vaccination. Data points represent individual animals; bars indicate mean values. Statistical significance was assessed using two-sided Mann–Whitney tests; only $P < 0.05$ is shown. The dotted line indicates the correlate of protection threshold (0.5 IU/mL).

**b,** Clinical scores following RABV challenge (RV437, 720 $TCID_{50}$), based on IACUC criteria including body weight, body condition, behavior and physical appearance. Lines represent individual animals.

**c,** Survival following RABV challenge at day 90 post-vaccination. Survival was analyzed by log-rank (Mantel–Cox) test. a-c, YF17D, n = 10; Rabipur n = 10; YF-RabG, n = 10.

**d-f**, Boostability of the antibody response at 4 months post-vaccination. YF-RabG-primed (n = 10, d) and Rabipur®-immunized (n = 6, e) hamsters were boosted with a 1/40 human dose of Rabipur®. Age-matched naïve hamsters (n = 6, f) receiving the same dose served as controls. The dotted line indicates the correlate of protection threshold (0.5 IU/mL).

#### EXTENDED DATA FIGURE 8

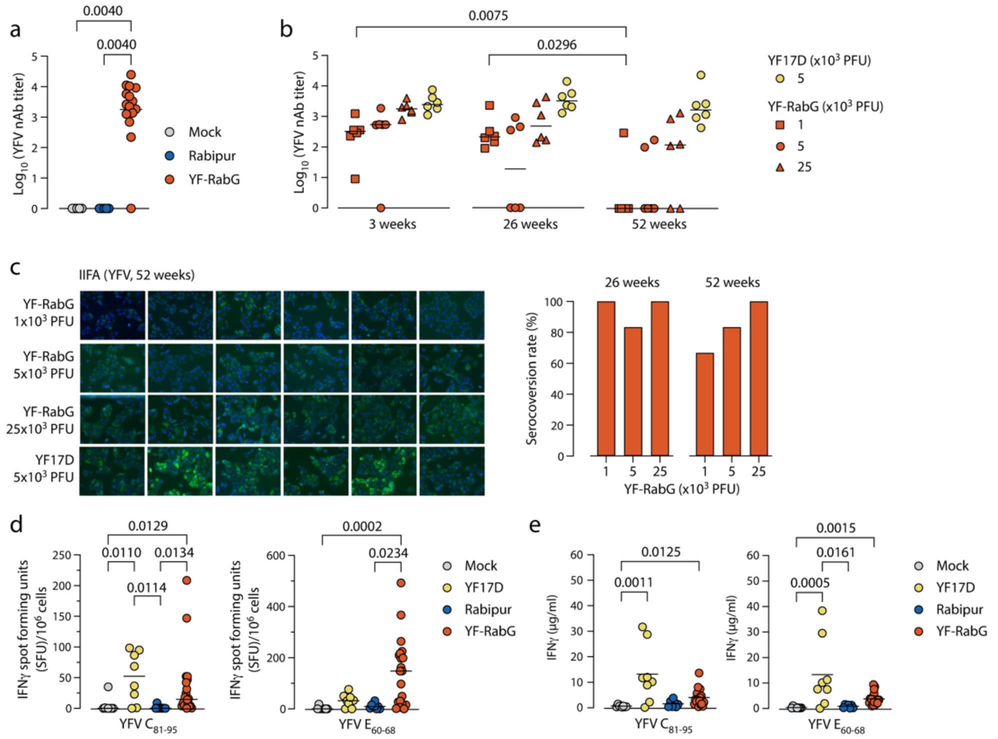


**Extended Data Fig. 8: Humoral and cellular immune responses to YFV elicited by YF-RabG.**

**a,** YFV-neutralizing antibody titers in *Ifnar*$^{-/-}$ mice at day 28 post-vaccination with Mock (n = 5), Rabipur® (n = 5) or YF-RabG (n = 15). Data points represent individual animals; bars indicate mean values. Statistical analysis was performed using Kruskal–Wallis test followed by Dunn's multiple-comparison test; only $P < 0.05$ are shown.

**b,** YFV-neutralizing antibody titers in hamsters at 3 and at 26 and 52 weeks post-vaccination with indicated doses of YF-RabG (1,000, 5,000 or 25,000 PFU; n = 6 per group) or YF17D (5,000 PFU; n = 6). Data points represent individual animals; bars indicate mean values. Statistical analysis was performed using two-way ANOVA with Tukey's multiple-comparison test; only P < 0.05 are shown.

**c**, Representative immunofluorescence images of fixed, permeabilized YFV-infected cells incubated with immune sera from collected 52 weeks after first vaccination. Total anti-YFV IgG are visualized in green; Nuclei in blue. The columns correspond to individual animals. Rows indicate the vaccine dose. On the right, a bar graph with the seroconversion rate of binding IgG antibodies at 26 and 52 weeks post vaccination with different doses of YF-RabG (as indicated).

**d, e,** Number of IFN-γ-secreting cells (d) and concentration of IFN-γ in culture supernatants (e) following *ex vivo* stimulation of splenocytes with the YFV-derived MHC class I-restricted envelope peptide $E_{60–68}$ or MHC class II-restricted capsid peptide $C_{81–95}$, respectively. Splenocytes were isolated from BALB/c mice vaccinated with YF-RabG ($2x10^3$ PFU; n = 20 in d; n = 19 in e), YF17D ($2x10^3$ PFU; n = 8) or Rabipur (1/40 human dose, administered on days 0 and 7; n = 8 in d; n = 7 in e), or from mock-vaccinated mice (medium containing 2% FBS; n = 8). Dots represent individual animals; bars indicate means. Statistical significance was assessed using Dunn's multiple-comparisons test; only $P < 0.05$ is shown.

**Supplementary Table 1**. List of antibodies used for flow cytometry.

| Antigen | Fluorophore | Working dilution | Clone | Cat. No. | Company |
|---|---|---|---|---|---|
| CD45 | BUV496 | 1:80 | D058-1283 | 741185 | BD |
| HLA-DR | BUV805 | 1:333 | G46-6 | 748338 | BD |
| CD20 | V450 | 1:167 | L27 | 655872 | BD |
| CD21 | BUV563 | 1:333 | B-Ly4 | 741362 | BD |
| CD27 | BUV395 | 1:100 | M-T271 | 740291 | BD |
| IgD | FITC | 1:40 | goat | 2030-02 | SBA |
| CD3 | BV650 | 1:100 | SP34.2 | 563916 | BD |
| CD4 | PE-Cy7 | 1:800 | L200 | 560644 | BD |
| CD8 | V500 | 1:167 | SK1 | 344732 | Biolegend |
| CD28 | BV785 | 1:80 | CD28.2 | 302950 | Biolegend |
| CD95 | BV711 | 1:167 | DX2 | 305644 | Biolegend |
| CD127 | PE-Cy5 | 1:40 | eBioRDR5 | 15-1278-42 | Invitrogen |
| CD25 | BUV737 | 1:40 | 2A3 | 612806 | BD |
| CD14 | ECD | 1:80 | RM052 | B92391 | Beckman Coulter |
| CD16 | BV605 | 1:40 | 3G8 | 563172 | BD |
| CD11c | Alexa700 | 1:20 | 3.9 | 56-0116-42 | Invitrogen |
| CD1c | APC | 1:20 | L161 | 17-0015-42 | Invitrogen |
| CD123 | PerCP-Cy5.5 | 1:100 | 7G3 | 558714 | BD |
| CD40 | BV750 | 1:100 | 5c3 | 746948 | BD |
| CD69 | APC-Cy7 | 1:80 | FN50 | 557756 | BD |
| CD71 | BV480 | 1:40 | L01.1 | 746247 | BD |
| CD80 | BUV661 | 1:40 | L307-4 | 741647 | BD |
| CD83 | BV421 | 1:20 | HB15e | 305324 | Biolegend |
| CD86 | Alexa647 | 1:67 | IT2.2 | 305416 | Biolegend |
| CD163 | PE | 1:20 | GHI/61 | 556018 | BD |
